# Effects of fire and fire-induced changes in soil properties on post-burn soil respiration

**DOI:** 10.1101/2024.04.23.590763

**Authors:** Dana B. Johnson, Kara M. Yedinak, Benjamin N. Sulman, Timothy D. Berry, Kelsey Kruger, Thea Whitman

## Abstract

**Background:** Boreal forests cover vast areas of land in the northern hemisphere and store large amounts of carbon (C) both aboveground and belowground. Wildfires, which are a primary ecosystem disturbance of boreal forests, affect soil C via combustion and transformation of organic matter during the fire itself, and via changes in plant growth and microbial activity post-fire. Wildfire regimes in many areas of the boreal forests of North America are shifting towards more frequent and severe fires driven by changing climate. As wildfire regimes shift and the effects of fire on belowground microbial community composition are becoming clearer, there is a need to link fire-induced changes in soil properties to changes in microbial functions such as respiration in order to better predict the impact of future fires on C cycling.

**Results:** We used laboratory burns to simulate boreal crown fires on both organic-rich and sandy soil cores collected from Wood Buffalo National Park, Alberta, Canada, to measure the effects of burning on soil properties including pH, total C, and total nitrogen (N). We used 70-day soil incubations and two-pool exponential decay models to characterize the impacts of burning and its resulting changes in soil properties on soil respiration. Laboratory burns successfully captured a range of soil temperatures that were realistic for natural wildfire events. We found that burning increased pH and caused small decreases in C:N in organic soil. Overall, respiration per gram total (post-burn) C in burned soil cores was 16% lower than in corresponding unburned control cores, indicating that soil C lost during a burn may be partially offset by burn-induced decreases in respiration rates. Simultaneously, burning altered how remaining C cycled, causing an increase in the proportion of C represented in the modelled slow-cycling vs. fast-cycling C pool as well as an increase in fast-cycling C decomposition rates.

**Conclusions:** Together, our findings imply that C storage in boreal forests following wildfires will be driven by the combination of C losses during the fire itself as well as fire-induced changes to the soil C pool that modulate post-fire respiration rates. Moving forward, we will pair these results with soil microbial community data to understand how fire-induced changes in microbial community composition may influence respiration.

## Introduction

Boreal forests cover 18.9 million km^2^ in the northern hemisphere (Brandt et al. 2013) and store vast amounts of carbon (C) above and belowground (Bradshaw and Warkentin 2015), making them an important global reservoir of C. The boreal zone is classified as the area of land at high northern latitudes that circumnavigates the Arctic and contains forests of cold-tolerant plant species and alpine areas (Brandt et al. 2013). Long, cold winters and short, cool summers lead to low rates of decomposition, supporting C accumulation (DeLuca and Boisvenue 2012). These C stores are affected by fire; both wildfire and prescribed fire used as a land management tool by indigenous groups have been an integral part of the boreal forests of Canada and Alaska for hundreds to thousands of years (MacDonald et al. 1991, Prince et al. 2018, Christianson et al. 2022). Understanding the impacts of wildfire on soil C stores over time is particularly important for predicting how the ecosystem services of boreal forests will change under future climate regimes.

During a wildfire, above and belowground organic matter (OM) can volatilize and combust, resulting in large C and nitrogen (N) emissions from the burned area – an estimated 51 Tg C year^-1^ was emitted during wildfires in the boreal forest of North America between 1997 and 2016 (Zhao et al. 2021). Combustion of the soil organic horizon is a large contributor to these C emissions. For example, soil C losses of 3.0 kg C m^-2^ were observed following a large wildfire in the boreal forest of the Northwestern Territories, Canada (Walker et al. 2018). In another study, 2.3 kg C m^-2^ was lost from the O horizon and mineral soil of a Douglas-fir forest during a wildfire in southwestern Oregon, USA (Bormann et al. 2008). However, the impacts of wildfire on boreal forest soil C stocks extend beyond loss of C during the wildfire itself. Fire also has immediate effects on soil properties, plant communities, and microbial community composition with long-term consequences for C cycling. Previous work using 2-pool exponential decay models showed a decrease in size (as fraction of total) of fast cycling C pools and a corresponding increase in decay rates of this pool following burning (Johnson et al. 2023), suggesting that fire-induced changes to the soil environment impact the cycling of the C that remains post-fire.

Changes to the soil environment are often driven by the temperatures experienced at the soil surface during a wildfire. Temperatures at the soil surface during a crown fire can reach hundreds of degrees Celsius (DeBano 1991, Neary et al. 1999). High temperatures can drive heat-induced denaturation of organic acids and the creation of base cation-rich ash, causing an increase in soil pH, which can persist for years following fire (Certini 2005, Úbeda et al. 2005). Understanding the effects of wildfires on soil pH in both the short and long-term is particularly important for predicting soil C cycling, as pH can influence bacterial community composition (Lauber et al. 2009, Rousk et al. 2010). We previously found an average pH increase of 2.7 units following burning of sandy acidic soil samples and organic-rich soil samples from the boreal forest (Johnson et al. 2023). However, the relationship between fire and soil pH is not straightforward. Under conditions of soil heating at moderate temperatures, pH has also been observed to decrease, which may be due to the volatilization and downward movement of organic acids from overlying horizons that then condense upon reaching the cooler underlying soil (Luo 2023). Additional research is needed to untangle the relationship between heat-induced changes in soil pH and post-fire soil C cycling.

Another way in which immediate effects of wildfire may influence longer-term soil C pools is via the incomplete combustion of OM and resulting creation of pyrogenic OM (Knicker 2011). Pyrogenic OM is characterized by polycyclic aromatic structures that lend it biological and chemical stability (Ohlson et al. 2009), generally reducing its bioavailability for microbes. Previously, we found that changes in C composition and related changes in degradation rates played a large role in driving depressed post-burn soil respiration rates (Johnson et al. 2023). Therefore, it is possible that burn conditions that favor pyrogenic OM production may have larger impacts on respiration rates (normalized by total remaining C) than burn conditions that cause more complete combustion of OM.

Partial or complete combustion of soil surface horizons and associated OM also kills microbes, immediately altering microbial community composition (Johnson et al. 2023, Certini et al. 2021). Heat-induced changes to soil properties such as those detailed above continue to affect soil microbial community composition months to years following fire, with potential consequences for C mineralization. For example, in a meta-analysis of microbial biomass following prescribed fires and wildfires, Dooley and Treseder (2012) documented a correlation between decreased microbial biomass and decreased soil respiration in boreal forest soils following wildfire. However, previously, we concluded that soil C mineralization rates are likely not constrained by loss of potential functions within microbial communities – *i.e.*, after burning, microbial communities could still readily degrade OM (Johnson et al. 2023). However, this previous study used homogenized soil subsamples to measure soil respiration, thereby obscuring the constraints of spatial accessibility of C to soil microbes.

In addition, burn-induced effects on soil properties are also influenced by fire intensity. This is increasingly important, given our new awareness of changing boreal forest wildfire regimes driven by climate change. Here we follow Keeley et al. (2009) using fire intensity to denote heat dosage and fire severity to mean ecosystem response, specifically loss of above and belowground organic matter. Wildfire regimes in the North American boreal forest are currently undergoing a shift towards more severe fires (Flannigan et al. 2000, Kasischke and Turetsky 2006, De Groot et al. 2013) as well as larger and more numerous fires (Natural Resources Canada 2019). With climate change and increasing temperatures at northern latitudes, fire severity is predicted to continue to increase and fire return intervals to shorten in the coming century (Balshi et al. 2009, Flannigan et al. 2009, Hanes et al. 2018), which may have large implications for these ecosystems. Fire severity influences the amount of time required for ecosystem recovery (Holden et al. 2016, Dove et al. 2020), and increasing fire severity in boreal forest ecosystems has been linked to significant decreases in post-burn soil respiration (Holden et al. 2016, Kelly et al. 2021). Understanding the effects of fire of varying severity on soil C cycling across a range of soil types is important for predicting boreal forest carbon fluxes under future climate conditions. Our objectives for this study were to quantify the effects of burning and burn duration on total C, total N, soil pH, and soil CO_2_ efflux in contrasting soil types typical of the Canadian boreal forest. We used a laboratory mass loss calorimeter to simulate boreal crown fire exposures characteristic of boreal crown fires on both intact organic-rich and sandy soil cores collected from Wood Buffalo National Park, Alberta, Canada, to evaluate the effects of burning. We conducted 70-day soil incubations and fitted two-pool exponential decay models to respiration data to measure the impact of burning and burn-induced changes to soil properties on soil respiration. Burning and incubating intact soils cores allowed us to account for spatial patterns in combustion and microbial mortality within a soil. This is important because soil temperature during a wildfire rapidly attenuates with depth due to the insulating properties of soil (Beadle 1940), and thus both mass microbial mortality and OM combustion are generally confined to the uppermost soil horizons. We hypothesized that (1) burning would cause an increase in soil pH and a decrease in total soil C and N, with larger changes following longer burns, (2) pH would decline towards (but not reach) pre-burn levels within months of burning, and (3) soil respiration would decline. In line with previous work, we expected to see a decrease in the fractional size of the fast-cycling C pool and corresponding increase in decay rate with burning, and an increase in the fractional size of the slow-cycling C pool and a corresponding decrease in decay rate with burning.

## Methods

### Study area

Soil cores were collected from 12 sites across Wood Buffalo National Park, located at the border of Alberta and the Northwest Territories, Canada, over a 5-day period in June 2022 (Supplementary Table 1). Average annual rainfall for this region is 300-400 mm, and average summer and winter temperatures range from 10 to 22 _J and -24 to -14 _J respectively (Alberta Climate Information Service 2018). Sites representing two dominant vegetation types – jack pine (*Pinus banksiana* Lamb.) and spruce (*Picea mariana* (Mill.) and/or *Picea glauca* (Moench) Voss) – were randomly selected from areas of forest unburned for greater than 30 years (Beaudoin et al. 2017, Natural Resources Canada 2022). Sites were located between 0.1 and 0.5 km from roads and > 0.5 km from other sampling sites. Upon arriving, we confirmed the dominant tree species. Soils underlying *Picea* spp.-dominated sites tended to have thick (>10 cm deep) organic horizons (Supplementary Table 2) consisting of living and dead sphagnum moss and are classified as Dystric Histosols (Food and Agriculture Organization of the United Nations 2003). Soils underlying *Pinus banksiana*-dominated sites tended to have thinner (0.5-5 cm thick) O horizons underlain by sandy mineral soil and are classified as Eutric Gleysols (Food and Agriculture Organization of the United Nations 2003).

### Sampling methods

At each site, sixteen soil cores 6 cm in diameter and 10 cm in height were collected across a 2 m radius circle using a soil core sampler with clear PETG plastic core liners (Giddings Machine Company, Windsor, CO, USA). Cores were labeled and capped with a perforated top cap to prevent development of anoxic conditions. We deconstructed three cores within 8 hours of sampling to measure O and mineral horizon thickness, bulk density, and field moisture. All collected soil was transported to Madison, WI within ten days of sampling and allowed to air dry in a dark room for 6 weeks to simulate drought conditions (Alberta Climate Information Service 2018). One core from each site was destructively sampled for a soil texture analysis using a physical analysis hydrometer at the UW-Madison Soil and Forage Lab (Madison, WI).

### Laboratory Burns

To simulate typical fires for this region, four cores from each site were exposed to 60 kW m^-2^ in a Mass Loss Calorimeter (Fire Testing Technology Limited, West Sussex, UK) for 120 second (Supplementary Figure 1). We chose this method for delivering the heat dose treatment over others (such as heating in an oven or water bath) to more closely mimic the spatial and temporal heat dose delivery of a natural wildfire. During the heat dose treatment, a spark ignitor located above the top of the core (Supplementary Figure 1) was used to trigger combustion. Following the treatment, each core was removed from the mass loss calorimeter, allowed to continue smoldering in a fume hood, and left to cool to room temperature. Heat flux and duration were chosen to represent wildland fire propagation (Silvani and Morandini 2009) for a mid to high-range crown fire (Frankman et al. 2012, Thompson et al. 2015) (Figure 1). Another four cores from each site were exposed to the same heat flux for 30 seconds. This duration was chosen to represent soil partially shielded from a flaming front by debris such as downed logs. Four additional cores from each site were not burned (designated “control” cores).

**Figure 1.**
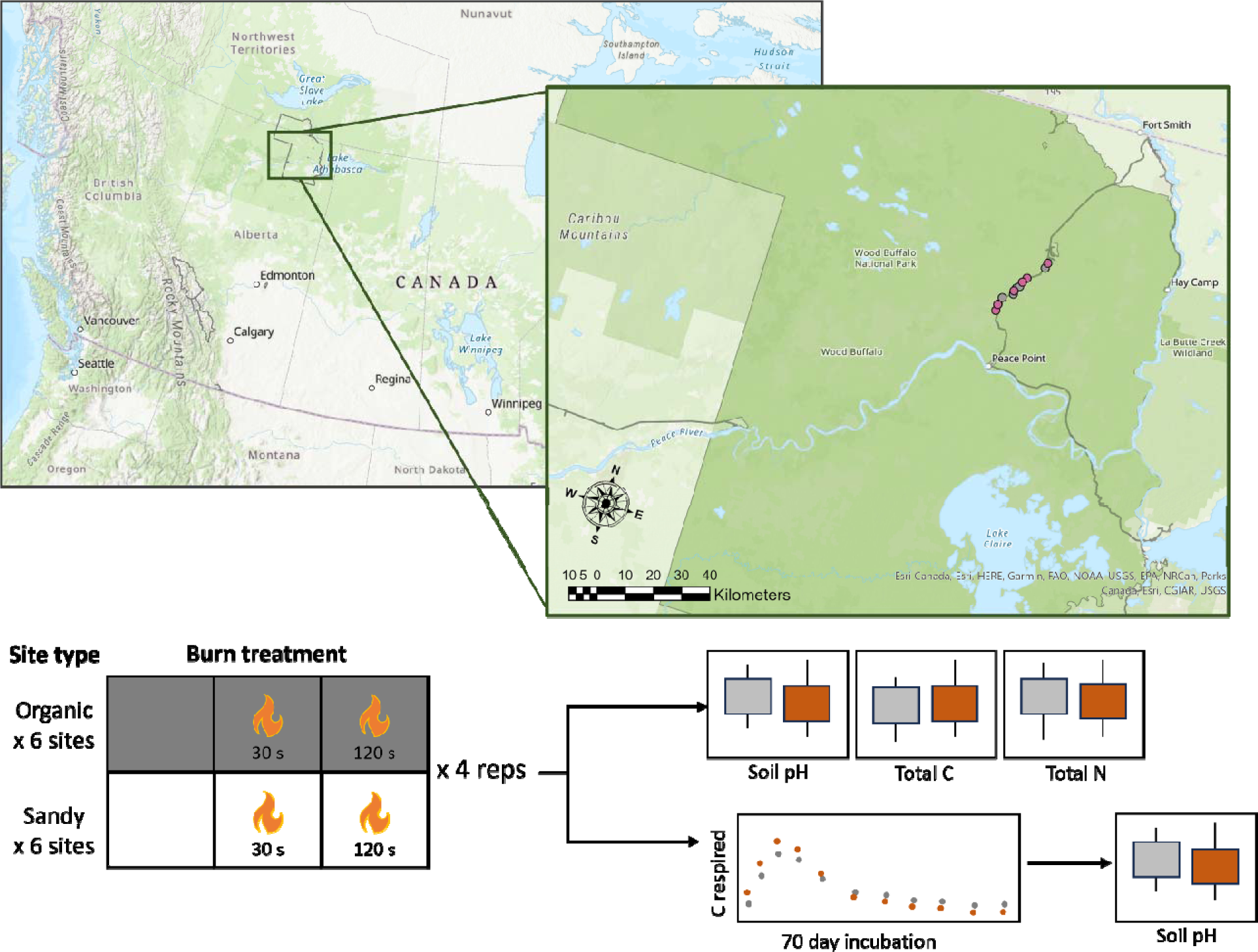
Experimental design. Soil cores were collected from 12 sites (pink and grey circles denote sites where Pinus banksiana and Picea spp., respectively, are the dominant vegetation) within Wood Buffalo National Park (top right), which is located at the northern border of Alberta, CA (top left). Soil cores were subjected to laboratory burns (unburned, 30 s heat dose treatment duration, and 120 s heat dose treatment duration). Soil CO_2_ efflux from burned and unburned soil was measured for 70 days.

Beaded 20-gauge Type-K thermocouples (GG-K-20-SLE, Omega) inserted horizontally at the base of the O horizon (or at 5 cm depth in cases where cores consisted of solely O horizon) and 1 cm above the core base of the core tracked temperature during the burns and for a minimum of 5 hours afterwards. To account for both the duration and degree of heating, we used a weighted measure of the cumulative time above 25 °C, designated “degree hours” (DH). DH were calculated by multiplying the timestep by each temperature data point (minus 25 °C) and summing the result across the portion of the temperature profile (Figure 2) above 25 °C. Twenty-four hours after the burn, mass loss and the depth of O horizon combustion was recorded. Sets of cores representing the three treatments (120 s burn duration, 30 s burn duration, and unburned control) from each site were incubated together for 48 hours, 24 days, 49 days, and 70 days.

**Figure 2.**
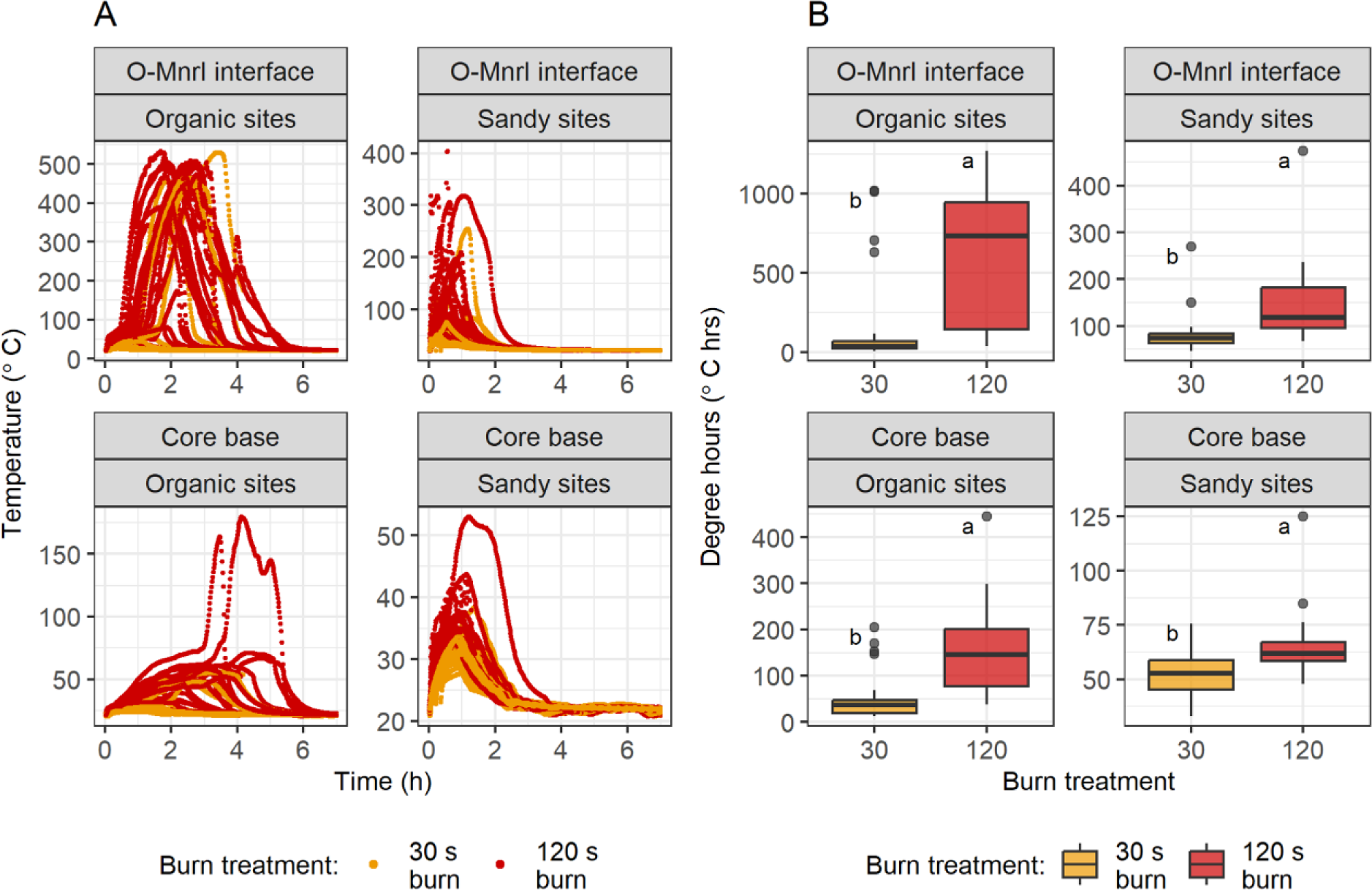
(A) Thermocouple temperatures and (B) degree hours (above 25 °C) in soil cores at the organic-mineral interface (top panels) and base (bottom panels) of cores from organic sites (left) and sandy sites (right) during and following the heat dose treatment (n=24). Heat dose treatments were initiated at t=0. Temperatures greater than 100 °C suggest the presence of flaming combustion and are not a quantitative measure of soil matrix. Different letters represent statistically significant differences in treatments based on ANOVA and Tukey’s HSD (p<0.05). For boxplots, the central horizontal line indicates the median, the upper and lower bounds of the box indicate the inter-quartile range (IQR), the upper and lower whiskers reach the largest or smallest values within a maximum of 1.5 * IQR, and data beyond the whiskers are indicated as individual points.

### Post-burn soil analyses

Water was added to intact soil cores to bring moisture to 65% of field capacity, and cores were placed in 475 mL Mason jars to incubate in the dark at room temperature. The cores were divided into four sets each containing three cores (unburned, 30 s burn, 120 s burn) from each site. The first set was destructively sampled 48 hours post-burn and subsamples of O and mineral horizon soil were collected for pH measurements (Braus and Whitman 2021) and C and N analysis (Flash EA 1112 CN Automatic Elemental Analyzer (Thermo Finnigan, Milan, Italy)). The second and third sets of cores were destructively sampled for pH measurements 24 days and 49 days post-burn, respectively. The final set of cores was used to track soil respiration over the course of a 70-day incubation, after which they were also destructively sampled for pH measurements. These cores were placed in 475 mL Mason jars fitted with gas-tight lines connected to an automated sample analyzer (“multiplexer”). The multiplexer automatically samples and quantifies the amount of CO_2_ in the jar headspace via a Picarro cavity ring-down spectrometer (Berry et al. 2021). Headspace CO_2_ concentrations were measured at least once per 24-hour period. To characterize microbial C mineralization, we fit two-pool exponential decay models for each sample to this flux data and estimated respiration rate constants and fractional sizes of fast and slow C pools using (Eq. 1):

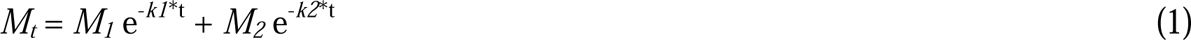

where *M_t_* is the total C pool, *M_1_* and *M_2_* are the fast-cycling and slow-cycling C pools, respectively, *k_1_* and *k_2_* are the respiration rate constants for the fast and slow C pools, respectively, and *t* is time. This model was fit using the *nls.lm* function in the *minpack.lm* package (Elzhov et al. 2016) in R.

### Statistical analyses

All analyses were performed using R statistical software version 4.3.1 (R Core Team 2023) and model results plotted using R package *ggplot2* (Wickham 2016). We used Wilcoxon signed rank tests to analyze differences in soil temperature during and following the heat dose treatments. We used ANOVA models and Tukey’s HSD to assess the impact of burning and heat dose treatment duration on soil pH, total C, total N, and C:N ratios. We used Kruskal-Wallis and paired Wilcox tests to analyze differences in respiration and decay model coefficients between burned and unburned soil.

## Results

### The effect of burning on soil temperature varied with soil type and heat dose duration

The two heat dose treatment durations (30 s and 120 s) created a range of soil temperatures and mass losses (Figure 2; Supplementary Figures 2-4; Table 1) representative of soil temperatures during a wildfire. Maximum temperatures at 5 cm depth in organic soil cores following the 120 s heat dose treatments (mean=336.8 ± 197.0 °C) were significantly higher than maximum temperatures following the 30 s heat dose treatments (mean=109.7 ± 168.7 °C) (Wilcoxon signed rank test, p=3.7x10^-5^). Maximum temperatures at the base of the O horizon in sandy soil cores following the 120 s heat dose treatments (mean=163.4 ± 94.7 °C) were significantly higher than maximum temperatures following the 30 s heat dose treatments (mean=67.2 ± 45.1 °C) (Wilcoxon signed rank test, p=9.1x10^-5^).

**Table 1.**
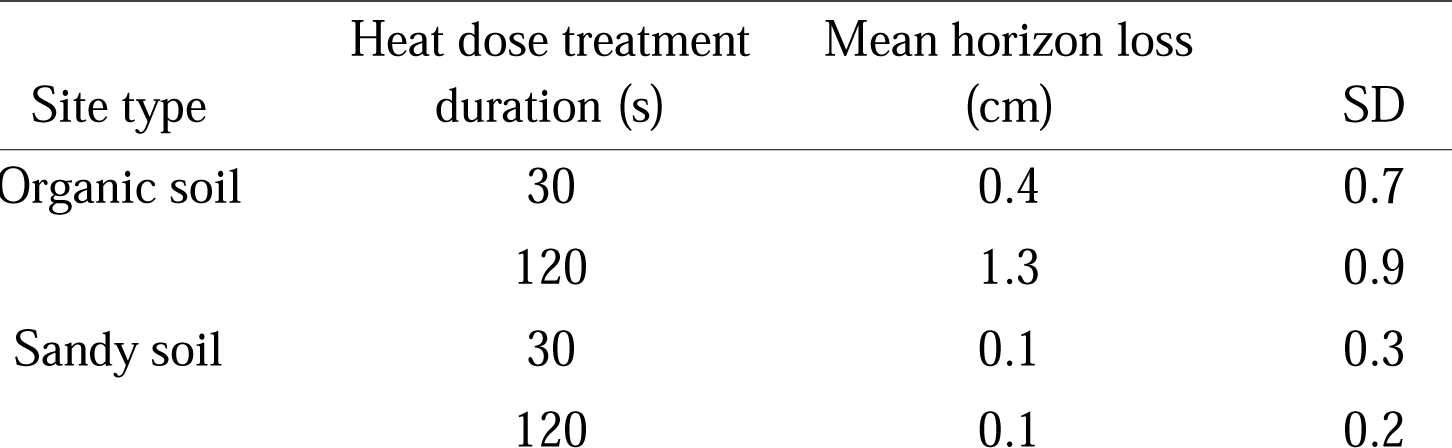
Horizon depth loss following heat dose treatments.

Temperatures at the base of the organic soil cores in 120 s burns reached a maximum of 180.1 °C (mean=56.9 ± 38.0 °C). Temperatures at the base of the sandy soil cores did not exceed 53.1 °C. All cores returned to room temperature within 6.3 hours.

Due to the complex nature of fire and the heterogeneity of intact soil cores, maximum temperatures reached following the heat dose treatment varied greatly between cores both within the same site and across sites (Supplementary Figure 3).

We recorded significantly higher DH at the upper thermocouple following the 120 s heat dose treatments in both organic (mean=626.1 ± 435.8 °C·hr) and sandy soil cores (mean=148.3 ± 86.3 °C·hr) compared to the 30 s heat dose treatments (organic, mean=172.8 ± 315.1 °C·hr; ANOVA, p=3.7x10^-5^; sandy, mean=83.8 ± 44.3 °C·hr; ANOVA, p=0.002).

### Soil pH increases with higher temperatures and longer heat exposures

Burning caused an immediate increase in soil pH (Figure 3). Following the 120 s heat dose treatment, soil pH was significantly higher in organic soil (mean=7.7±0.7), sandy soil O (mean=6.1±1.1), and mineral horizons (mean=5.0±0.5) compared to unburned controls (organic soil, mean=6.6±0.7; ANOVA, p=0.01; sandy soil O horizon, mean=4.6±0.2; ANOVA, p=0.01; sandy soil mineral horizon, mean=4.8±0.5; ANOVA, p=0.03). There was only weak evidence that the 30 s burn duration treatment had a positive effect on soil pH in organic soil (mean=7.3±0.6; ANOVA, p=0.1). There was no significant change in pH following the 30 second heat exposure in sandy soil (O horizon, mean=5.0±0.4; ANOVA, p=0.6; mineral horizon, mean=4.9±0.5; ANOVA, p=0.4). The interaction between site type (organic *vs.* sandy) and heat dose treatment duration was not significant.

**Figure 3.**
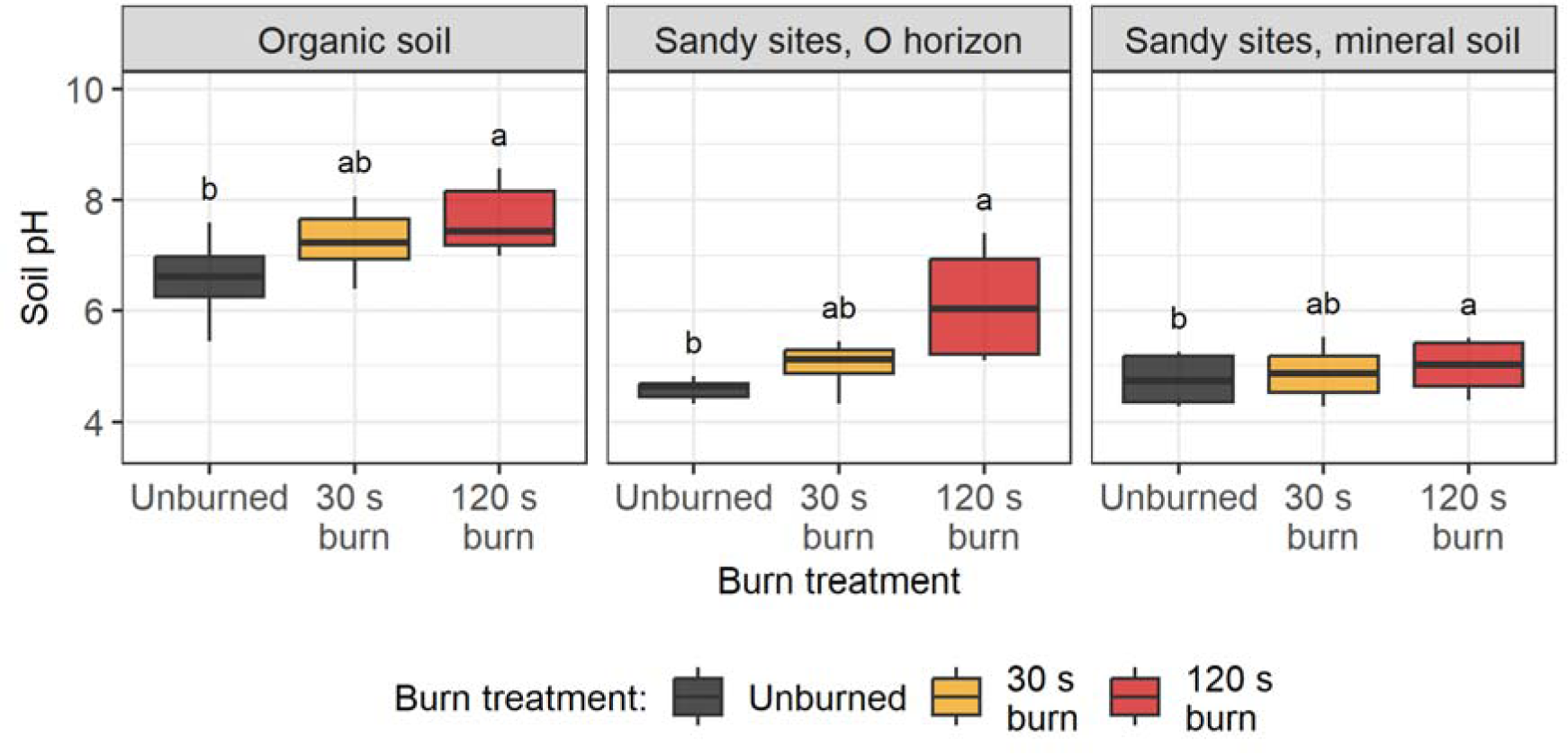
Soil pH two days post-burn in in organic soil (left panel), sandy O horizons (middle panel), and sandy mineral soil (right panel). Different letters represent statistically significant differences in treatments based on ANOVA and Tukey’s HSD (p<0.05). The central horizontal line indicates the median, the upper and lower bounds of the box indicate the inter-quartile range (IQR), the upper and lower whiskers reach the largest or smallest values within a maximum of 1.5 * IQR, and data beyond the whiskers are indicated as individual points.

Soil pH was significantly positively correlated with DH (p=0.01, R*_adi._*^2^=0.73) in organic and sandy soil (interaction term between site type and soil pH, p=2.1x10^-5^) (Supplementary Figure 5).

Soil pH increased over the course of the 70-day incubation in control cores for organic cores (2 days post-burn, mean=6.6 ± 0.7; 70 days post-burn, mean=7.8±0.9; ANOVA, p=2.6e-02) and for the organic horizon of sandy cores (2 days post-burn, mean=4.6 ± 0.2; 70 days post-burn, mean=5.3±0.5; ANOVA, p=1.4e-02). There was no change in pH of mineral horizons of control samples during the incubation. For the burned cores, there were significant differences in pH over the course of the incubation only in the organic horizons of sandy cores exposed to the 30 second heat treatment (2 days post-burn, mean=5.0 ± 0.2; 70 days post-burn, mean=5.7±0.4; ANOVA, p=5.6e-02) (Supplementary Figure 6).

### Soil C:N ratios decrease with burning

Despite losses in total stocks through combustion, burning did not significantly change soil total C or N concentrations relative to unburned soil (Supplementary Figures 7 and 8); however, burning did affect C:N ratios. Following the 120 s heat dose treatment, soil C:N ratios were significantly lower in organic soil (mean=26.3±6.9) compared to unburned controls (organic soil, mean=30.7±6.4; ANOVA, p=0.04) (Figure 4). There was no difference in C:N ratios between burned and unburned sandy soil horizons. The interaction between site type (organic *vs*. sandy) and heat dose treatment duration in ANOVA model for C:N ratios was not significant. There was a modest decrease in C:N ratios with increasing DH in cores from both organic site and O horizons of sandy sites (Supplementary Figure 9) (linear model, p=0.04, R*_adj_*^.2^=0.10).

**Figure 4.**
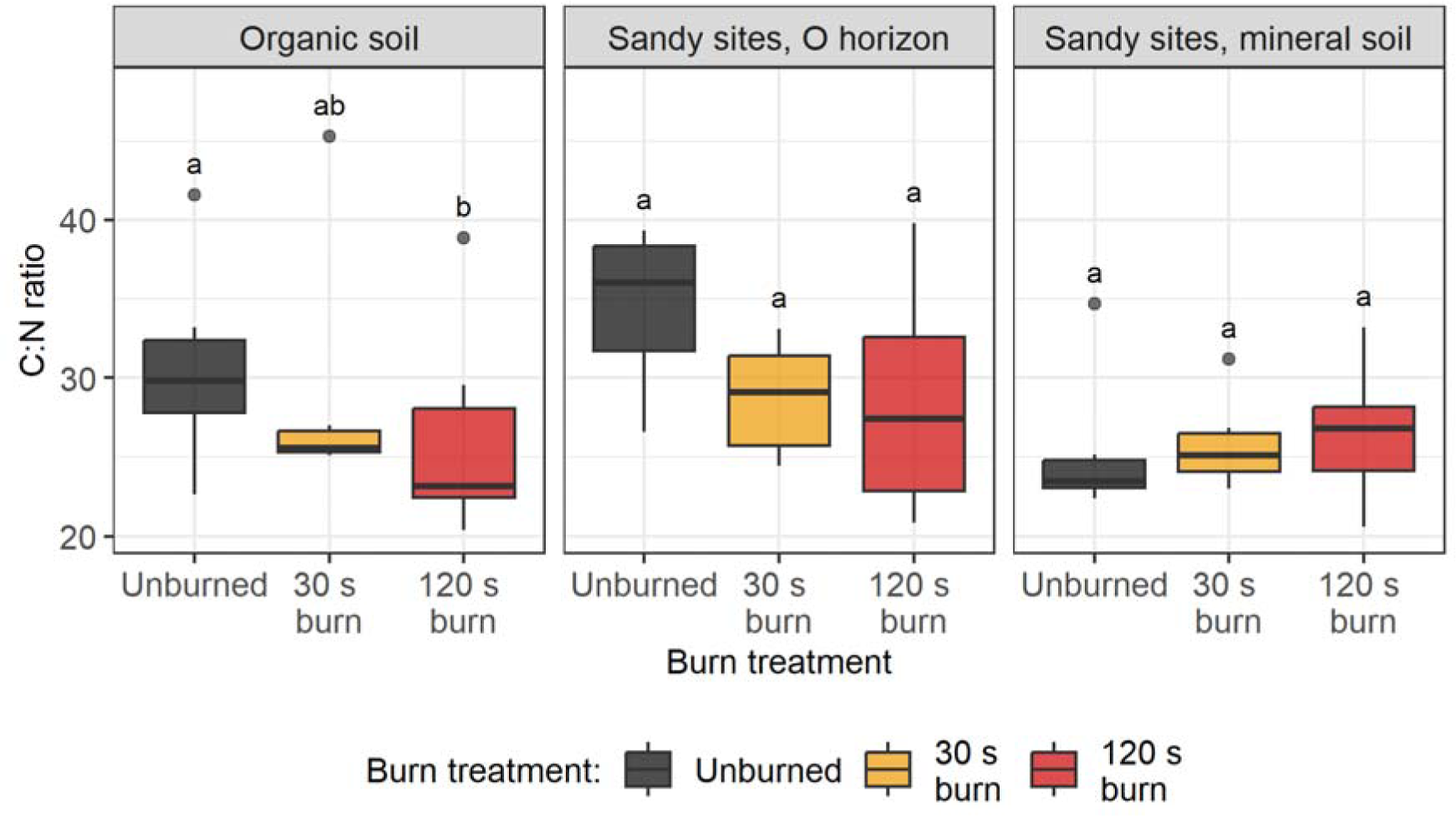
Soil C:N ratio with heat dose treatment for different soil types and horizons. Different letters represent statistically significant differences in treatments based on ANOVA and Tukey’s HSD (p<0.05). The central horizontal line indicates the median, the upper and lower bounds of the box indicate the inter-quartile range (IQR), the upper and lower whiskers reach the largest or smallest values within a maximum of 1.5 * IQR, and data beyond the whiskers are indicated as individual points.

Due to difficulties in disentangling mass loss from water loss during a burn event, we estimated C and N mass pre and post-burn by multiplying pre-burn soil volume by dry bulk density estimates and soil C concentration estimates from unburned control cores (Supplementary Figure 8). Following the 120 s burn duration treatment, average C mass loss from the organic and sandy soil was estimated to be 1.2±1.4 g and 0.2±0.3 g, respectively. Average N mass loss from the organic and sandy soil was estimated to be 0.042±0.05 g and 0.007±0.012 g, respectively.

### Burning decreased respiration rates

Total C respired as a fraction of C remaining post-burn over the course of the 70-day incubation in cores from both organic and sandy sites was significantly lower in burned soil (organic, 120 s burn duration, mean=2.1±0.5; sandy, 120 s burn duration, mean=2.2±1.6) compared to unburned organic (organic, mean=4.4±0.4; ANOVA, p=0.0002) and sandy cores (mean=5.4±2.7; ANOVA, p=0.05) (Figure 5, Supplementary Figure 10).

**Figure 5.**
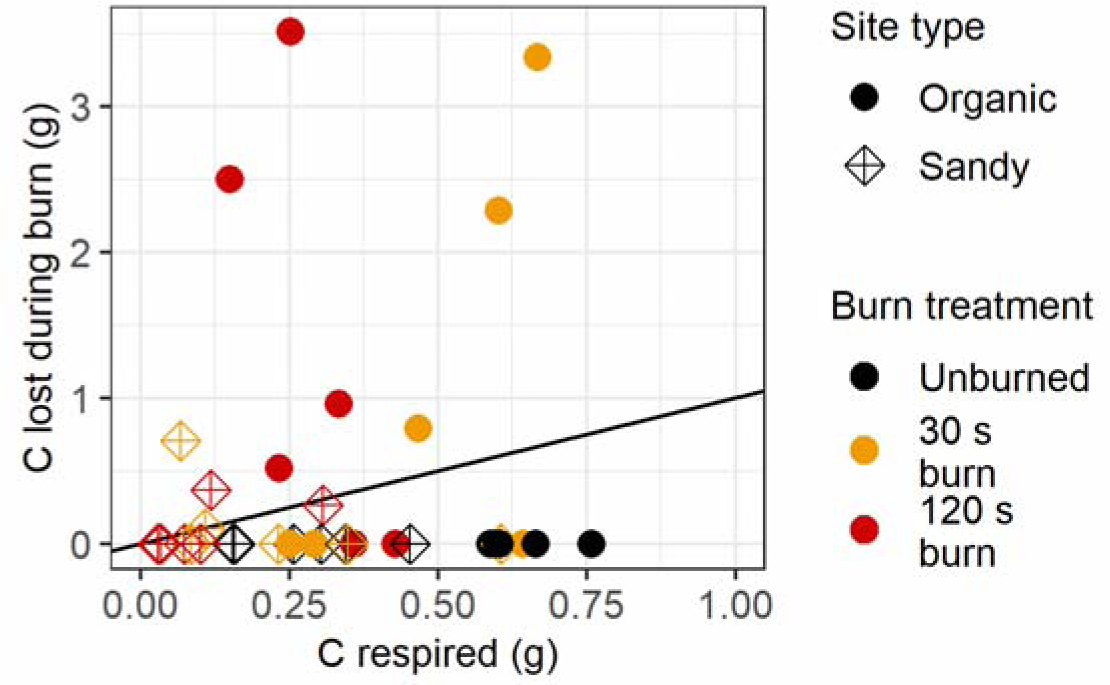
Mass C lost to burning vs. mass C respired during the 70 day incubation. The black line indicates a 1:1 relationship.

Two-pool decay models resulted in good fits to respiration data from the 70-day incubation (Supplementary Figure 11; R^2^ > 0.998). In both organic and sandy soil cores, burning increased fast-cycling C pool decomposition rates and increased the proportion of C held in the slow-cycling C pool (Figure 6). Higher fast-cycling C pool decomposition rates (*k_1_*) were found following the 120 s burn duration treatment in both organic (mean=0.1±0.1) and sandy cores (mean=0.1±0.1) compared to unburned organic (mean=3.8x10^-3^±2.7x10^-3^; Kruskal Wallis test, p=0.007) and sandy cores (mean=8.2x10^-3^±6.3x10^-3^; Kruskal Wallis test, p=0.003) (Figure 6).

**Figure 6.**
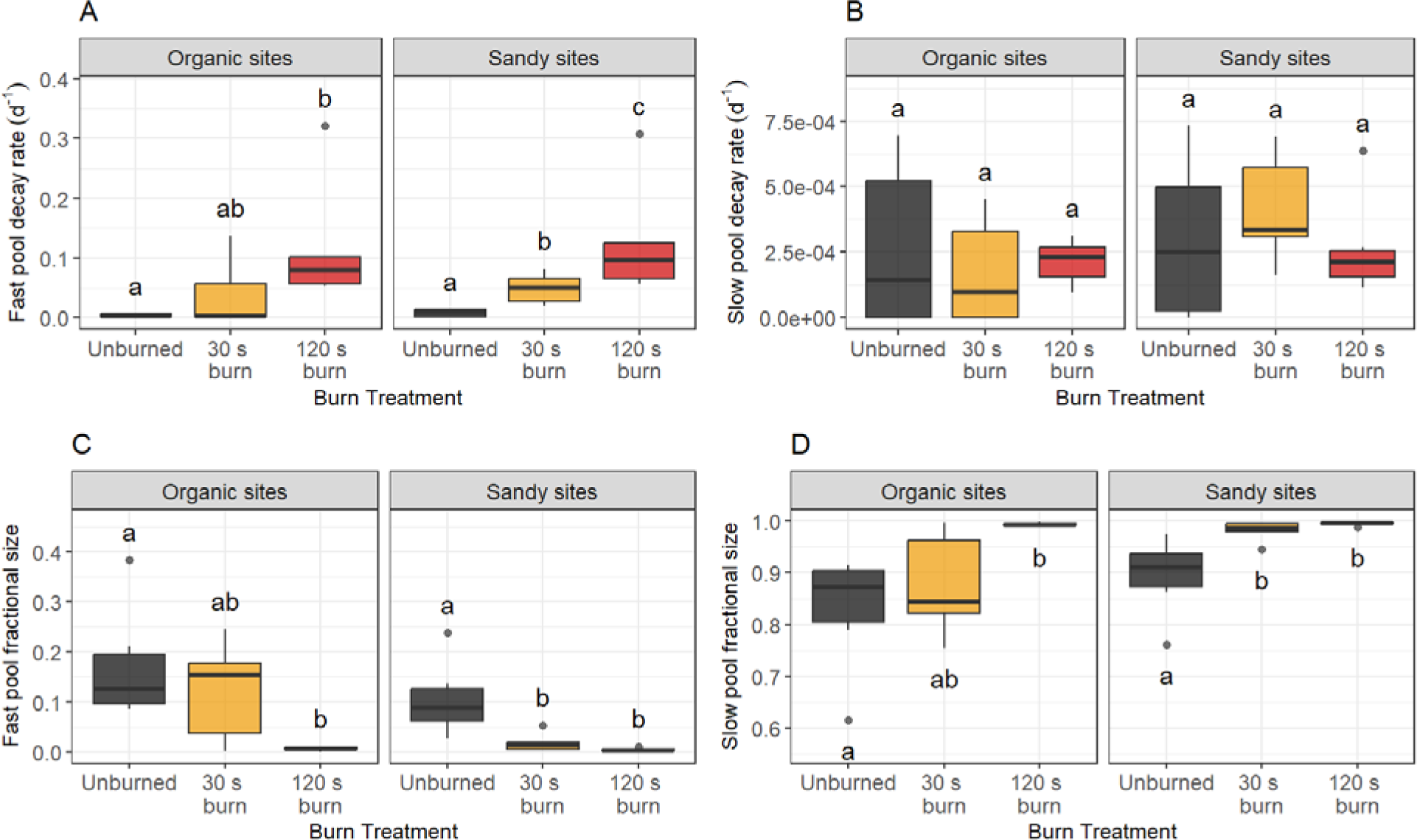
Coefficients for (A, B) decay rate and (C, D) fractional C pool size. Different letters represent statistically significant differences in treatments based on Kruskal Wallis test and Pairwise Wilcoxon test (p<0.05). The central horizontal line indicates the median, the upper and lower bounds of the box indicate the inter-quartile range (IQR), the upper and lower whiskers reach the largest or smallest values within a maximum of 1.5 * IQR, and data beyond the whiskers are indicated as individual points.

Based on respiration measurements following the 120 s burn duration treatment, the fraction of C held in the fast-cycling C pool (*M_1_*) decreased in both organic (mean=6.4x10^-3^±3.7x10^-3^) and sandy cores (mean=4.2x10^-3^±3.3x10^-3^) compared to unburned organic (mean=0.2±0.1; Kruskal Wallis test, p=0.007) and sandy soil cores (mean=0.1±0.1; Kruskal Wallis test, p=0.007) (Figure 6).

## Discussion

### Immediate effects of burning on soil differ with soil type and burn duration

Heat dose treatments of short and long durations resulted in a range of soil temperatures realistic to a natural wildfire event within a boreal forest. Temperatures at the soil surface during a crown fire can reach hundreds of degrees Celsius (DeBano 1991, Neary et al. 1999), although there is also a high degree of spatial heterogeneity for surface temperature and combustion (Lavoie and Mack 2012). Our two distinct durations used as heat dose treatments resulted in generally distinct temperature profiles, with longer duration heat dose treatments resulting in higher soil DH (Figure 2; Supplementary Figure 2), which allowed us to compare the effects of different heat dose treatment duration on soil properties and soil respiration.

High soil temperatures (mean=169 _J) were reached in the upper half of the cores, but temperature rapidly attenuated with depth, consistent with the expectation that direct burn effects will largely be confined to surface soil horizons. This is supported by previous work finding larger reductions in microbial biomass in surface (≤5 cm) soils than in subsoils (Pressler et al. 2019, Johnson et al. 2023). However, maximum temperature reached (Supplementary Figure 3) and duration of elevated temperature were different between cores from the two soil types. The higher temperatures and higher DH recorded in organic cores compared to sandy cores (Figure 2) were most likely driven by differences in the quantity of OM. The sandy cores were composed primarily of mineral soil, which is not susceptible to a large degree of combustion during a fire, and a thinner O horizon (1 – 4.5 cm in depth) at the surface (Supplementary Table 2).

In contrast, the organic cores were composed entirely of OM with no mineral soil. This OM was composed of partially decomposed *Sphagnum* moss with a low moisture content due to the drying period designed to represent drought conditions before the burn. While we may expect large C losses under future fire regimes, the effects of fire on soil can be highly variable – i.e., individual patches of soil may have significantly different heating and combustion during a wildfire. For example, numerous studies have documented high spatial heterogeneity in the degree of O horizon combustion during a single fire with areas of unburned O horizon existing adjacent to bare mineral soil (Kasischke and Johnstone, 2005; Lewis et al. 2011). Thus, in order to understand the impact of fire on soil C cycling, we must consider variability in fire intensity. The temperatures reached during burns in this experiment reflect this heterogeneity. While temperatures were generally higher in organic cores, this was not a universal rule. For example, following the 120 s burn duration treatment, temperatures greater than >300 °C were reached in 5 of 24 sandy cores, while only 7 of 24 organic cores failed to reach this threshold (Supplementary Figure 2). Additionally, there were a few instances in which a core exposed to the shorter burn duration (30 s) reached higher temperatures than corresponding cores from the same site that were exposed to the 120 s burn duration (Supplementary Figure 2). At the end of both the 30 s and 120 s heat dosing treatments, cores were removed from the mass loss calorimeter, but no steps were taken to stop ongoing combustion. As such, we attribute the cases of higher temperatures following 30 s burns relative to 120 s burns to continued combustion of OM after the completion of the heat dosing treatment.

### High temperatures and longer heat exposures increase soil pH

Despite large variation in temperature and DHs across cores, our hypothesis that burning would cause an increase in soil pH was generally supported by the 120 s burn duration treatments (Figure 3). While the shorter heat dose treatment did not have a large effect on soil pH, the positive correlation between DH and soil pH across all burned samples (Supplementary Figure 5) is also consistent with our expectations.

Fire-driven increases in soil pH are attributed to ash deposition and thermal modification of OM (Giovannini and Lucchesi 1997, Bodí et al. 2014, Araya et al. 2017), with higher soil temperatures resulting in larger pH increases up to a point (Araya et al. 2017, Bárcenas-Moreno et al. 2022). Over time, erosion and leaching remove ash from the soil profile, resulting in soil pH returning to pre-burn levels (Bodí et al. 2014, Matosziuk et al. 2020). Our study did not include leaching or erosion, which may explain why we did not observe pH declining towards pre-burn levels over the course of the 70-day incubation (Supplementary Figure 6). Another possible explanation is that the mechanisms driving post-fire pH recovery act over time periods longer than 70 days. The persistence of elevated pH post-fire has been documented years after fire in some field studies (Ulery et al. 1993, Dove et al. 2020), which suggests that any potential effects of pH on other properties, such as soil C cycling, could also be persistent.

The effects of fire on soil pH are further complicated by the myriad of other mechanisms influencing pH, such as active acidification by vegetation. The organic cores in this study had thick layers of *Sphagnum* spp. growing at the core surface at the time of sampling. *Sphagnum* has been shown to actively acidify its surrounding environment by releasing uronic acid into soil solution (Rydin et al. 2006). We observed an increase in soil pH in unburned organic soil and sandy soil O horizons over the course of the 70-day incubation (Supplementary Figure 6), which was unexpected. We believe that by incubating soil cores in the dark, we inhibited *Sphagnum*’s ability to engineer its external environment, which may have caused the observed increase in soil pH in unburned soil. In contrast, in the burned soils, since we killed the *Sphagnum*, this mechanism would not have been active, further explaining why we observed persistently elevated pH in burned soils. In mineral soil, we did not see a meaningful change in pH over the course of the 70-day incubation, which is also consistent with the previous explanation as the mineral horizons contained no *Sphagnum*. This highlights the role of vegetation recovery in driving post-fire recovery of soil function. *Sphagnum* has been observed to fully recover to pre-burn abundance within 10-50 years of fire in boreal peatlands and subalpine and alpine bogs, (Kuhry 1994, Clarke et al. 2015), although less time may be necessary for the acidifying effects of *Sphagnum* to occur.

### Burning decreased soil C:N despite small effects on total C and N concentrations

Our hypothesis that burning would cause a decrease in total soil C and N concentrations was not supported by our data (Supplementary Figure 7), despite estimated losses of up to 3.5 g C and 0.12 g N from burned cores (Supplementary Table 3). Combustion of OM at and near the soil surface removes C and N from the soil, and soil C is lost more quickly than N due to lower temperature thresholds for complete C volatilization relative to N (Araya et al. 2017). While this does not necessarily affect overall C and N concentration due to the relatively large pool of C and N deeper in the soil that is mostly untouched by burning, larger C than N losses may explain our findings of lower C:N ratios in organic soils following the 120 s heat dose treatment duration compared to unburned soil (Figure 4), despite the lack of significant differences in total C and total N concentrations (Supplementary Figure 7).

Recovery of C:N ratios to pre-fire levels will likely follow vegetation recovery and the resulting changes of plant inputs to the soil OM pool. Litter and belowground inputs to organic soils consist of *Picea* spp. branches, stems, and needles as well as material from ericaceous plants and *Sphagnum* moss. While we did not measure C:N ratios at the end of the 70-day soil incubation, we would not expect significant changes in these ratios due to lack of plant inputs to soil in this study. However, microbial activity and differential mineralization of C *vs*. N could potentially drive changes in C:N over time.

In contrast to the organic soils, in mineral soils, we did not observe significant differences between burned and unburned C:N ratios (or soil C concentrations and N concentrations). Temperatures required to volatilize mineral soil components are much higher than those required for OM combustion (Neary et al. 1999); therefore, we would not expect to see significant mineral soil mass loss. Loss of C and N from mineral soil via OM combustion may occur at the top of the mineral soil horizon where hottest temperatures are reached; however, temperatures rapidly attenuate with depth (Figure 2), rendering high C and N loss at depth unlikely. Even relatively large losses of C and N from the top of horizon could be diluted beyond detection when mixed with underlying soil.

### Burning and burn-induced changes to soil properties decrease respiration

Our hypothesis that burning and burn-induced changes to the soil environment would cause declines in soil respiration was supported by our data. Post-burn soil CO_2_ efflux as a fraction of total post-burn C from both organic and sandy soils exposed to the 120 s burn duration treatment was lower compared to unburned soil (Supplementary Figures 10 and 11). This observation may be explained by a variety of factors, each of which we will discuss.

First, burning may affect soil respiration via direct effects on soil microbial communities. For example, heat-induced microbial mortality driven by high soil temperatures can lead to a decrease in microbial biomass following burning (Dooley and Treseder 2012, Pingree and Kobziar 2019). However, soil temperature rapidly attenuates with depth in the soil due to the insulating properties of soil (Beadle 1940) and, thus, mass microbial mortality (and OM combustion) is expected to be largely confined to the uppermost soil horizons. This is supported by lower subsoil temperatures than surface temperatures (Figure 2) and by our previous work finding a large decrease post-burn in DNA and RNA concentrations in O horizon soil but not in underlying mineral soil after an identical 120s heat treatment in cores from the same region (Johnson et al. 2023), indicating that decreased microbial community size may play a relatively minor role in shaping post-fire C cycling.

Numerous studies have documented correlations between decreased microbial biomass and decreased soil respiration following fire (Dooley and Treseder 2012, Holden et al. 2016); however, if microbial mortality imposed a large limit on soil respiration, in our study we would expect to see decreased decay rates for both fast and slow-cycling C pools in two-pool decay models. Instead, our results show an increase in fast-cycling C pool decomposition rate, suggesting that the ability of microbial communities to cycle C has not been seriously impaired by burning (Figure 6). Although the increase in the fast pool decay rate following burning is not likely to exert a strong influence on overall C cycling rates, given the small proportion of C held in the fast C pool, these results are in line with our previous findings of increased fast-cycling C decay rates and no impairment of total mineralization following burning (Johnson et al. 2023).

Thus, although fire decreases microbial biomass, decreased soil respiration post-fire is likely often driven by other factors, such as burning-induced changes to soil properties, including pH and C:N ratios, resulting in an unfavorable environment for C degradation.

Soil pH may influence post-fire respiration rates by altering availability of organic C to microbes. Organic C, which sorbs more readily to mineral surfaces at low pH, is released from mineral surfaces and becomes more accessible to microbial degradation as pH increases (Kupka and Gruba 2022). Soil pH is also positively correlated with enzymatic potential for oxidizing persistent soil OM (Sinsabaugh et al. 2008), which is particularly relevant in post-fire boreal forest soils where microbes encounter thermally-altered OM. Given this, we would expect increasing pH to result in increased soil respiration rates. Indeed in some instances, increasing pH in acidic soils to circumneutral levels has been shown to increase soil respiration rates resulting in increased soil C losses (Malik et al. 2018). However, in this study, we did not observe significant positive correlations between pH and C mineralization. Instead, we found weak evidence for a negative correlation between soil pH and total C respired (Supplementary Figure 12; linear model, p=0.04, R*_adj._*^2^=0.10), suggesting that, following fire, any effects of pH towards increasing C mineralization rates are small or offset by fire-induced changes to other soil properties.

Similar to expected pH effects, in N-limited boreal forests, decreasing soil C:N would be expected to alleviate N limitation leading to increased soil respiration, which is, again, opposite to our findings of decreased respiration rates post-fire despite decreased soil C:N (in organic soils). Using C:N ratios as a proxy for substrate availability for microbes admittedly relies on an overly simplistic conceptualization of C compounds post-fire, which brings us to a third factor that may be driving post-fire respiration rates – C remaining after a burn may be less accessible to surviving microbes.

Although we did not measure C chemistry directly, all the soil cores experienced some degree of partial combustion based on the appearance of ash and charred material at the soil surface. As OM and soil begin to heat up, volatile compounds like water vaporize and organic molecules begin to break down. Complete combustion leaves behind inorganic material in the form of ash. Incomplete combustion results in large, disordered molecules and pyrogenic OM, with a higher degree of aromaticity than starting OM. With increasing temperatures of formation, pyrogenic OM contains more aromatic hydrocarbons and fewer aliphatic hydrocarbons (Wiedemeier et al. 2015), which may make this material less microbially accessible (Whitman et al. 2013). Amending soils with pyrogenic OM produced at temperatures > 300 _J has been shown to reduce soil respiration (Whitman et al. 2014, Santos et al. 2021, Adkins and Miesel 2021), through a variety of potential mechanisms.

The thermal degradation of organic compounds may also explain our findings of decreased C:N ratios with burning because, as previously noted, during heating, soil C is lost more quickly than N due to lower temperature thresholds for complete C volatilization relative to N (Araya et al. 2017). Thus, it is possible that depressed post-burn soil respiration rates in this study are driven by the alteration of OM to include more chemically recalcitrant molecules, despite a lower C:N. The larger proportion of soil C in burned soils (subjected to the 120 s burn duration treatment) held in the slow-cycling C pool relative to unburned soils (Figure 6) is also evidence for fire-induced changes in soil C chemistry. Interestingly, we do document a brief period 1-3 days after the initiation of the 70-day incubation during which respiration rates in burned soil increase sharply to levels equal to or greater than unburned soil respiration rates, and then decline below unburned soil respiration rates (Supplementary Figure 13). This pulse in respiration could be driven by a burn-induced release of nutrients, such as killed plant roots and microbial necromass, that is quickly depleted by microbial activity, leaving more recalcitrant forms of C behind. Future studies would be required to confirm the causes of this phenomenon. Additionally, we found no significant difference in the decay rates of the slow-cycling C pool in burned or unburned soil (Figure 6), suggesting that burning did not select for microbes with enhanced abilities to degrade fire-altered compounds.

The intensity and duration of fire-induced changes to soil properties and microbial function may all contribute to shaping post-fire C mineralization over different timescales. Immediately post-fire nutrient availability may be temporarily elevated due to combustion and the release of nutrients from microbial and root necromass. On the other hand, the combination of OM loss and decreased post-fire C mineralization rates due to changes in substrate quality, and hence C availability, may result in lower nutrient availability over some longer time.

### C losses to burning as compared to changes in short-term C losses from post-fire soil respiration

While it can be difficult to quantify loss of soil C via combustion during a natural wildfire, through the use of soil cores and laboratory burns, we were able to estimate burn-induced C losses (Figure 5). Overall, by 70 days post-burn, burned soils lost more C than unburned soil, but, for most cores, the bulk of this C was lost during the burn itself, not via post-fire soil respiration. In fact, post-fire soil respiration was depressed, in both absolute (per g initial pre-burn C) and relative (per g remaining C) terms. This raises the question of whether decreased respiration in burned soils relative to unburned soil may serve as a mitigating factor of C loss following fire. Given the timing of boreal forest burn seasons (generally late-June through August) and the relatively short growing season of circumboreal ecosystems, our finding of decreased respiration persisting 70 days post-fire indicates that effects of wildfire on soil respiration will play out over months to years.

The degree to which decreased post-burn respiration rates will offset C losses during a fire depends on the persistence of depressed respiration rates in burned soils. The results of this study suggest that post-fire respiration rates are largely driven by burn-induced changes to C accessibility for microbes; thus, a return to pre-burn respiration rates will likely depend on a combination of recovery of OM inputs to soil and decreasing reliance on thermally altered organic molecules over time. Recovery of soil OM inputs is dependent upon vegetation recovery. Post-fire recovery of aboveground C in living vegetation can take years to decades (Palviainen et al. 2020). As fresh OM inputs increase towards pre-burn levels, the proportion of C in the ‘fast-cycling C pool’ would be expected to increase. At the same time, thermally altered OM would not be increasing, so its role as a fraction of respired C would presumably diminish, although it is also the case that weathering of pyrogenic OM over time would be expected to increase its microbial accessibility, and, hence its mineralization (Zeba et al. 2022). In a review of post-fire boreal forest soil emissions, researchers found that total soil CO_2_ fluxes continue to increase for 10-30 years post-fire (Ribeiro-Kumara et al. 2020), which supports the idea that as vegetation recovery plays out, the composition of soil C will gradually shift towards more microbially available substrates and respiration will return to previous levels. At the same time, total soil C pools will increase in size, driven by the recovery of inputs to the soil, thus offsetting the increase in soil respiration, and resulting in a net increase in – i.e., recovery of – total soil C stocks.

Given that the heat dose treatments increased the proportional size of the slow cycling C pool (Figure 6), and, by definition, the slow cycling C pool has a lower decomposition rate than the fast cycling C pool (here, by three orders of magnitude), thermally altered OM will most likely persist in the soil compared to unburned OM. The post-burn increase in the proportion of C in the slow-cycling pool may indicate that soil C losses during wildfires from OM combustion may be offset in the long-term by the creation of more persistent soil C. To understand drivers of soil respiration years post-fire, more work will be required, likely drawing on modelling long-term effects of fire on soil C cycling.

## Conclusion

Burning of boreal forest soils altered the post-fire soil environment by removing material via combustion and volatilization, increasing soil pH, and, in organic soils, decreasing C:N ratios. Simultaneously, burning altered how remaining C cycles, causing an increase in the proportion of C represented in the slow-cycling *vs.* fast-cycling C pool as well as an increase in fast-cycling C decomposition rates. The combined effects of burning and burn-induced changes to the soil environment decrease soil respiration, and lower respiration rates in burned *vs.* unburned soil persisted for months post-burn. Persistence of depressed respiration rates implies that soil C lost during the burn may be partially offset by post-fire soil respiration over months to years. These effects were generally larger following longer duration heat dose treatments, which may be viewed as a proxy for more severe fire events. Future work is needed to explore the persistence of post-fire depressed soil respiration rates *in situ*.

## Declarations

### Ethics approval and consent to participate

Not applicable

### Consent for publication

Not applicable

### Availability of data and material

The datasets generated and analysed during the current study are available at ESS-DIVE under XXX [TBD]. Code to analyze data and produce figures is available at GitHub at XXX [TBD].

### Authors’ contributions

T.W., B.N.S., and D.B.J. conceptualized and designed the study. D.B.J., K.M.Y., B.N.S., and T.W. developed the methodology. D.B.J. and T.W. developed the software. D.B.J., K.M.Y., K.K., T.D.B., and T.W. conducted investigations. D.B.J. wrote the original draft. D.B.J. performed visualizations. T.W. supervised and administered the project and T.W. and B.N.S. acquired funding. All authors reviewed and edited the manuscript.

## Acknowledgements

We thank K. Bourne and E. Lazarcik at the USDA FS Forest Products Laboratory for assistance with burn simulations; D. Letourneau and M.A. Parisien at Natural Resources Canada for assistance with field work; J. Morin, T.J. Little, and other Wood Buffalo National Park staff for support in conducting this research (Permit WB-2022-41998).

