## Supplementary Information for "Effects of fire and fire-induced changes in soil properties on post-burn soil respiration"

Table of Contents	Page

**Supplementary Tables**

Supplementary Table 1. Site location and characteristics.	2

Supplementary Table 2. C and N concentrations of unburned soil.	3

Supplementary Table 3. Estimated mass (g) of soil C and N loss with burning.	3

**Supplementary Figures**

Supplementary Figure 1. Mass loss calorimeter to simulate fire effects on soil.	4

Supplementary Figure 2. Temperature in soil cores.	5

Supplementary Figure 3. Maximum temperatures.	6

Supplementary Figure 4. Photos of unburned and burned soil cores.	7

Supplementary Figure 5. Soil pH two days post-burn vs. degrees hours.	7

Supplementary Figure 6. Soil pH over the 70 day incubation.	8

Supplementary Figure 7. Soil C and N concentrations with heat dose treatment.	9

Supplementary Figure 8. Estimates of grams C and N with heat dose treatment.	10

Supplementary Figure 9. Soil C:N ratios two days post-burn with increasing DH.	11

Supplementary Figure 10. C respired over the entire 70 day incubation.	11

Supplementary Figure 11. Fraction of remaining post-burn total C.	12

Supplementary Figure 12. Soil pH versus C respired.	12

Supplementary Figure 13. Soil respiration rates for the first 10 days.	13

### Supplementary Tables

Supplementary Table 1. Site location and characteristics.

|  | Site ID | Latitude | Longitude | Soil texture class | Dominant tree species |
| --- | --- | --- | --- | --- | --- |
| 1 | 1 | -112.26 | 59.49 | Loamy Sand | Pinus banksiana |
| 3 | 2 | -112.41 | 59.40 | Sand | Pinus banksiana |
| 6 | 3 | -112.49 | 59.33 | Sand | Pinus banksiana |
| 8 | 5 | -112.39 | 59.41 | Sandy Loam | Pinus banksiana |
| 13 | 8 | -112.48 | 59.36 | Silt Loam | Pinus banksiana |
| 14 | 9 | -112.41 | 59.38 | Sand | Pinus banksiana |
| 7 | 4 | -112.49 | 59.31 | Organic | Picea spp. |
| 10 | 6 | -112.25 | 59.51 | Organic | Picea spp. |
| 11 | 7 | -112.36 | 59.44 | Organic | Picea spp. |
| 16 | 10 | -112.42 | 59.39 | Organic | Picea spp. |
| 17 | 11 | -112.38 | 59.43 | Organic | Picea spp. |
| 18 | 12 | -112.49 | 59.33 | Organic | Picea spp. |

Supplementary Table 2. C and N concentrations on unburned soil.

| Site ID | Soil horizon | Depth of horizon  sampled (cm) | C concentration (%) | N concentration (%) |
| --- | --- | --- | --- | --- |
| 4 | O | 10.0 | 38.06 | 1.68 |
| 6 | O | 10.0 | 48.35 | 1.16 |
| 7 | O | 10.0 | 26.30 | 0.97 |
| 10 | O | 10.0 | 42.46 | 1.43 |
| 11 | O | 10.0 | 42.78 | 1.29 |
| 12 | O | 10.0 | 40.32 | 1.34 |
| 1 | O | 2.0 | 15.65 | 0.40 |
| 1 | M | 8.0 | 1.26 | 0.05 |
| 2 | O | 2.0 | 32.63 | 0.91 |
| 2 | M | 8.0 | 2.08 | 0.06 |
| 3 | O | 2.0 | 14.07 | 0.36 |
| 3 | M | 8.0 | 1.16 | 0.05 |
| 5 | O | 2.5 | 22.23 | 0.61 |
| 5 | M | 7.5 | 1.19 | 0.05 |
| 8 | O | 1.4 | 8.36 | 0.28 |
| 8 | M | 8.6 | 1.12 | 0.05 |
| 9 | O | 3.0 | 20.73 | 0.78 |
| 9 | M | 7.0 | 2.07 | 0.09 |

Supplementary Table 3. Estimated mass (g) of soil C and N loss with burning.

|  | Burn duration treatment (s) | Carbon loss (g) | | | Nitrogen loss (g) | | |
| --- | --- | --- | --- | --- | --- | --- | --- |
|  |  | mean | sd | max | mean | sd | max |
| Organic soils | 30 | 1.07 | 1.42 | 3.33 | 0.04 | 0.04 | 0.10 |
|  | 120 | 1.25 | 1.44 | 3.51 | 0.04 | 0.05 | 0.12 |
| Sandy soils | 30 | 0.27 | 0.57 | 1.41 | 0.01 | 0.02 | 0.04 |
|  | 120 | 0.21 | 0.33 | 0.74 | 0.01 | 0.01 | 0.03 |

### Supplementary Figures

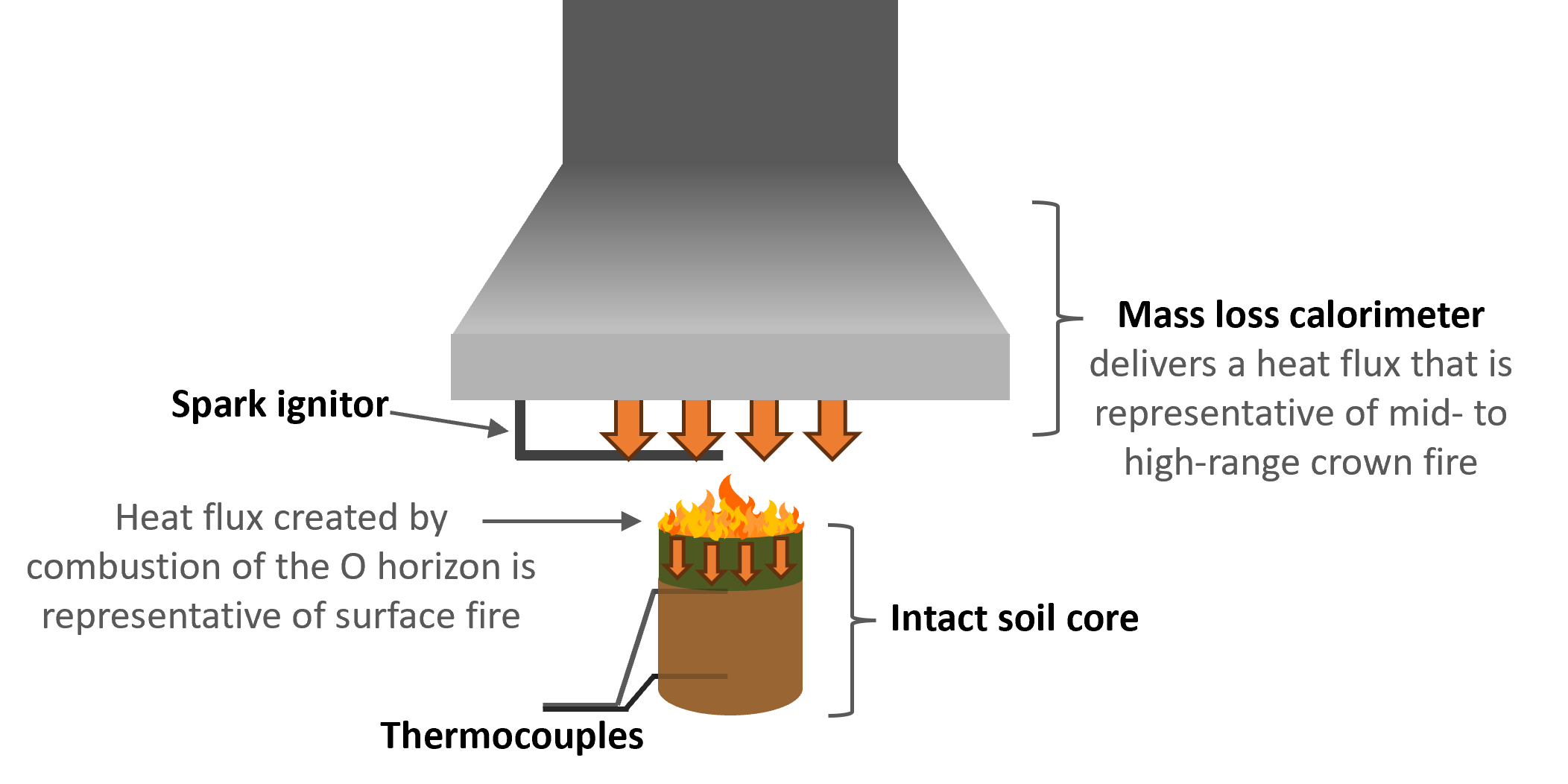

Supplementary Figure 1. Experimental use of a mass loss calorimeter to deliver a heat flux representative of a boreal forest crown fire.

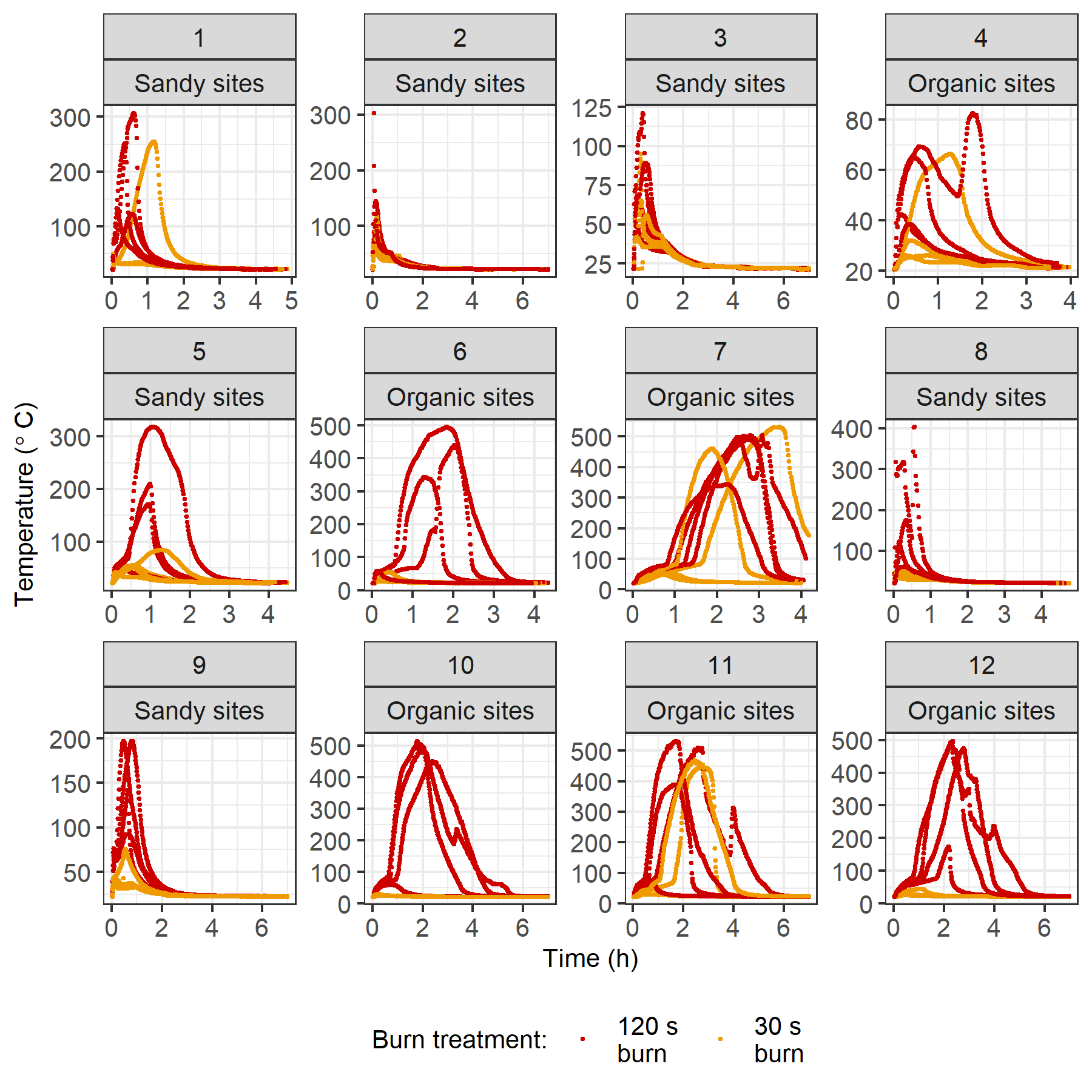

Supplementary Figure 2. Temperature in soil cores from organic sites and sandy sites during and following the heat dosing treatment as measured by thermocouples placed at the O-mineral interface.

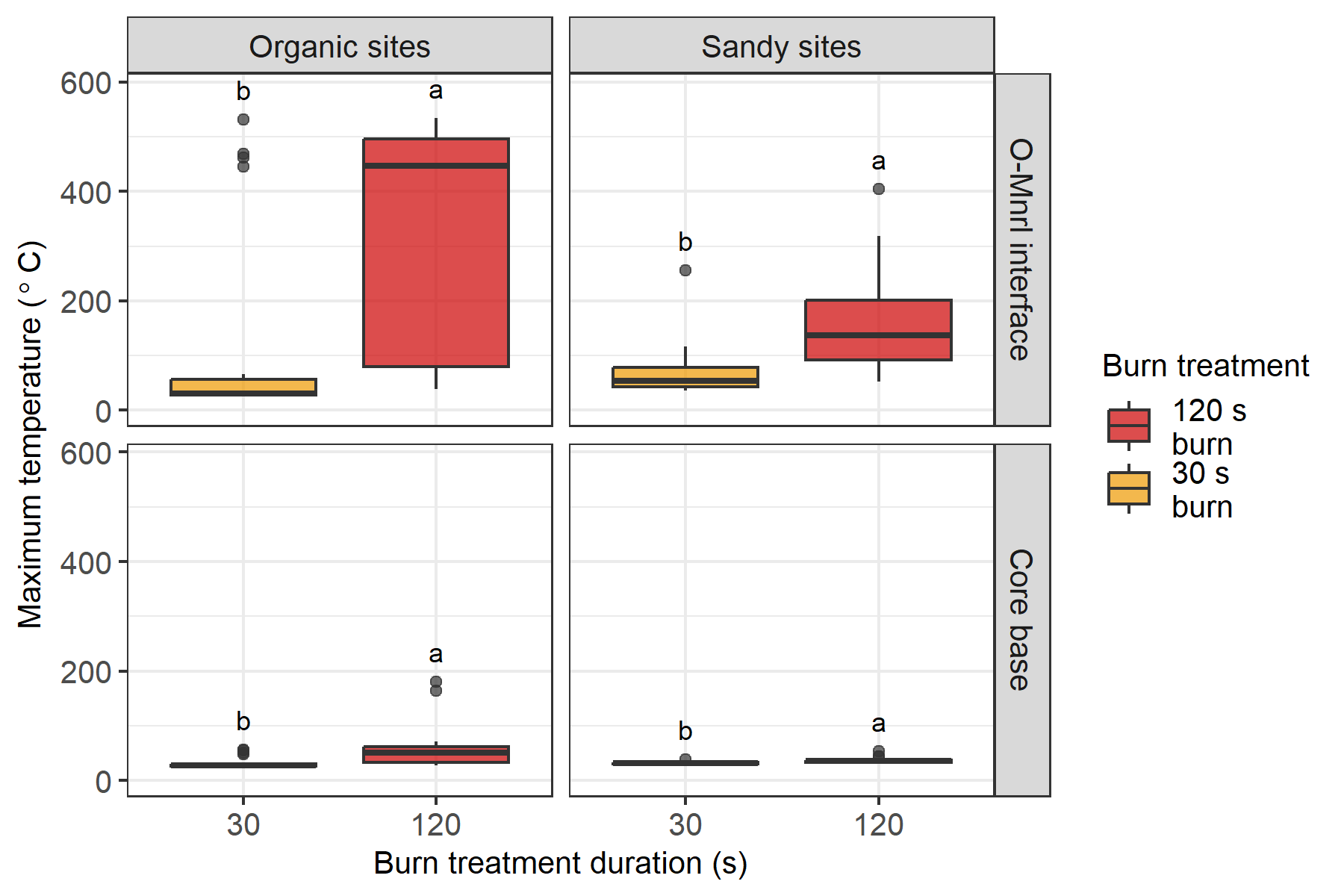

Supplementary Figure 3. Maximum temperatures reached in soil cores from organic sites (left) and sandy sites (right) during and following the heat dosing treatment as measured by thermocouples placed at the O-mineral interface (top panel) and core base (bottom panel). Different letters represent statistically significant differences in treatments based on Wilcoxon signed rank tests (p<0.05). The central horizontal line indicates the median, the upper and lower bounds of the box indicate the inter-quartile range (IQR), the upper and lower whiskers reach the largest or smallest values within a maximum of 1.5 * IQR, and data beyond the whiskers are indicated as individual points.

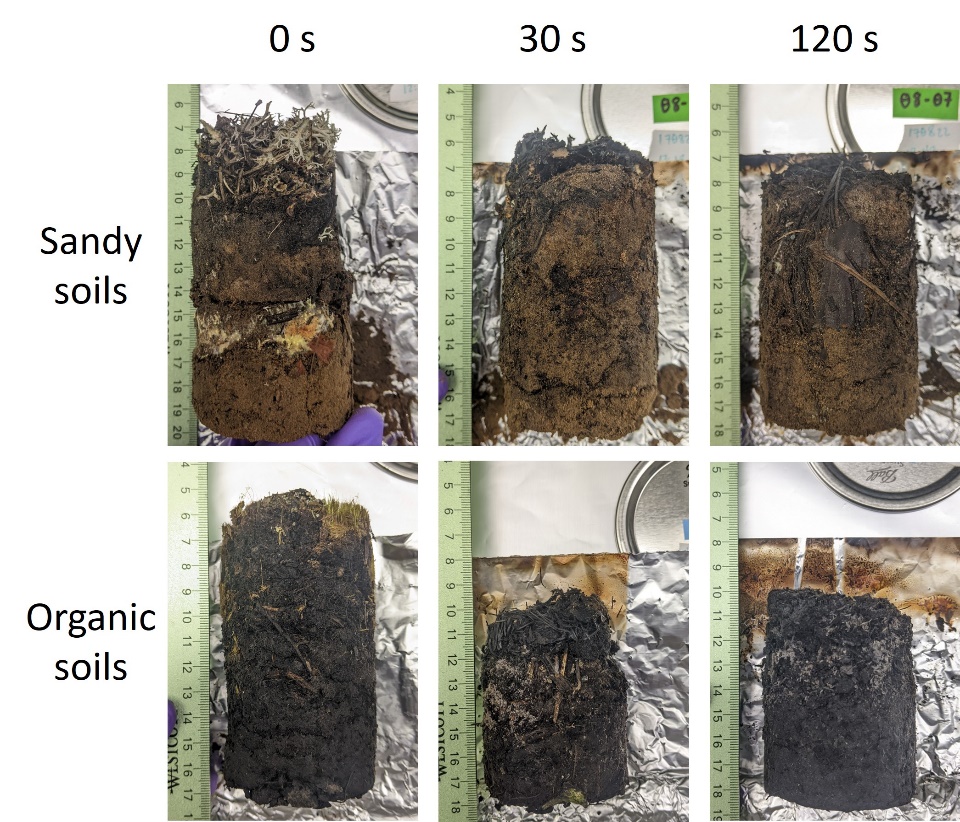

Supplementary Figure 4. Representative photos of unburned and burned soil cores 48 hours after the heat dose treatments.

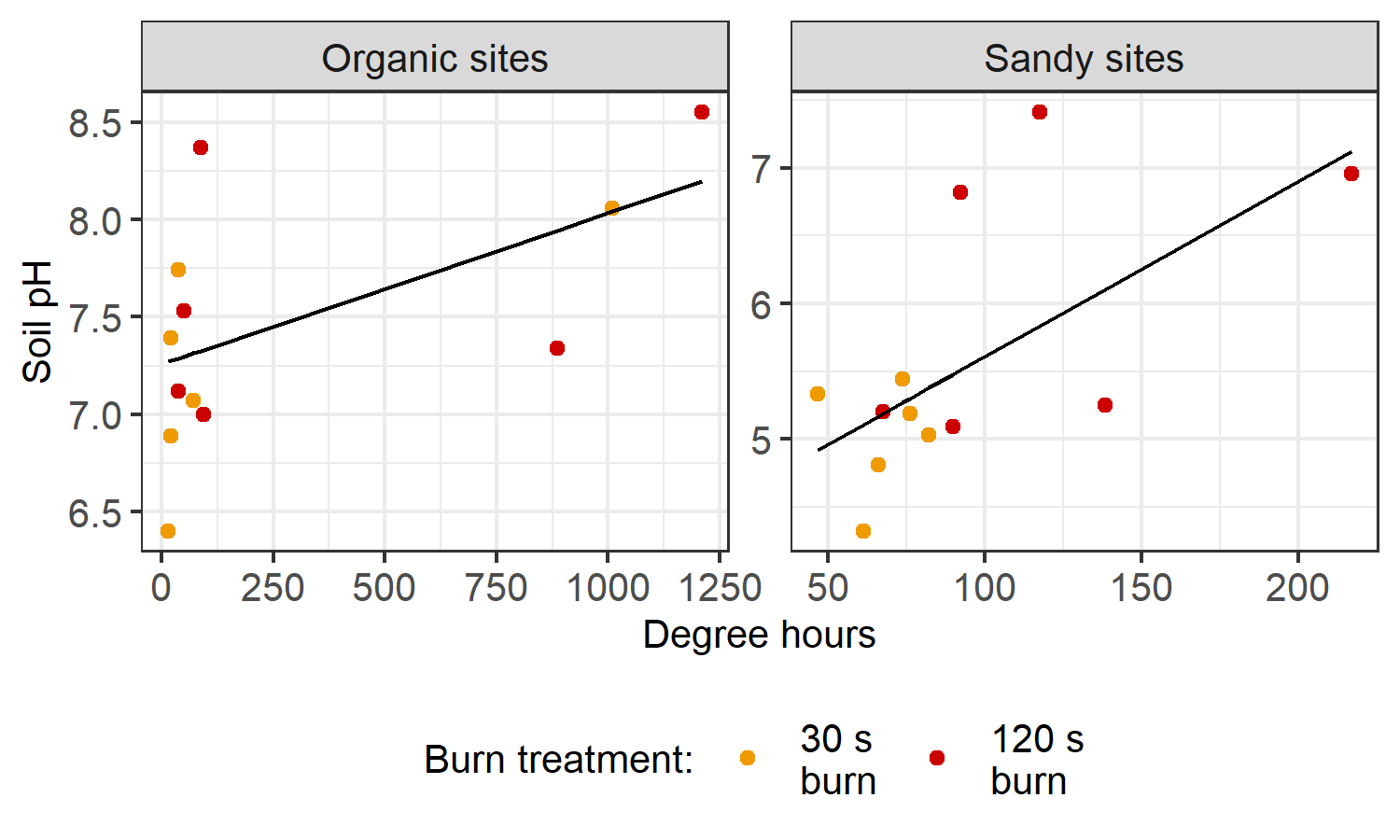

Supplementary Figure 5. Soil pH two days post-burn with increasing degree hours (DH) in O horizons of organic soils (left panel, y = 28.586 + -0.005x) and sandy soils (right panel, y = 34.469 + -0.061x) (n=12 soil horizon samples; p=0.01, $R_{adj.}^{2}$=0.73). DH were calculated based on thermocouple placed at O-mineral interface or at core midpoint. We used a linear model including an interaction term between site type and DH to test for a significant correlation between DH and soil pH (interaction term between site type and DH, p=2.1x10^-5^).

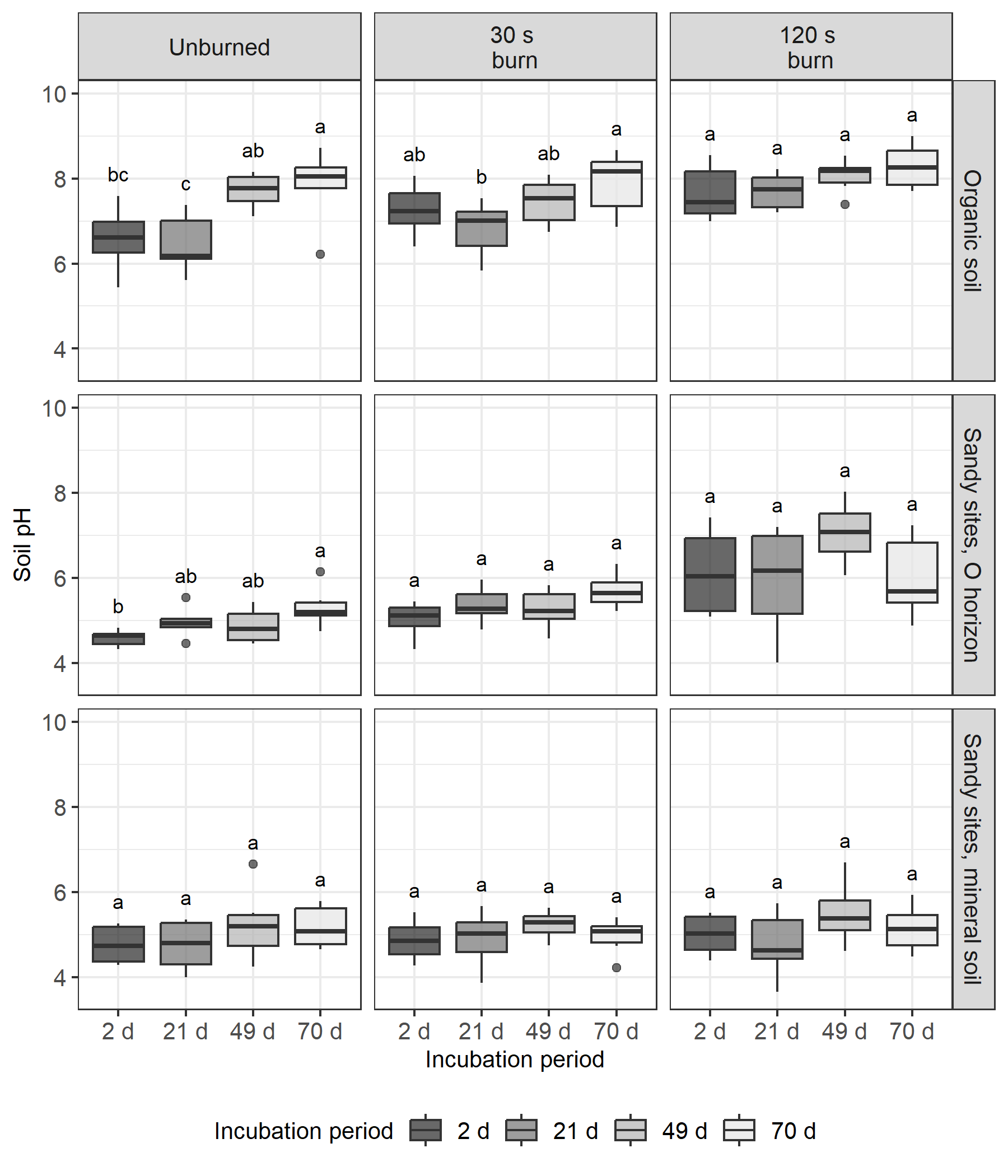

Supplementary Figure 6. Soil pH over the course of the 70-day incubation. Different letters represent statistically significant differences soil pH across timepoints based on ANOVA and Tukey’s HSD (p<0.05). The central horizontal line indicates the median, the upper and lower bounds of the box indicate the inter-quartile range (IQR), the upper and lower whiskers reach the largest or smallest values within a maximum of 1.5 * IQR, and data beyond the whiskers are indicated as individual points.

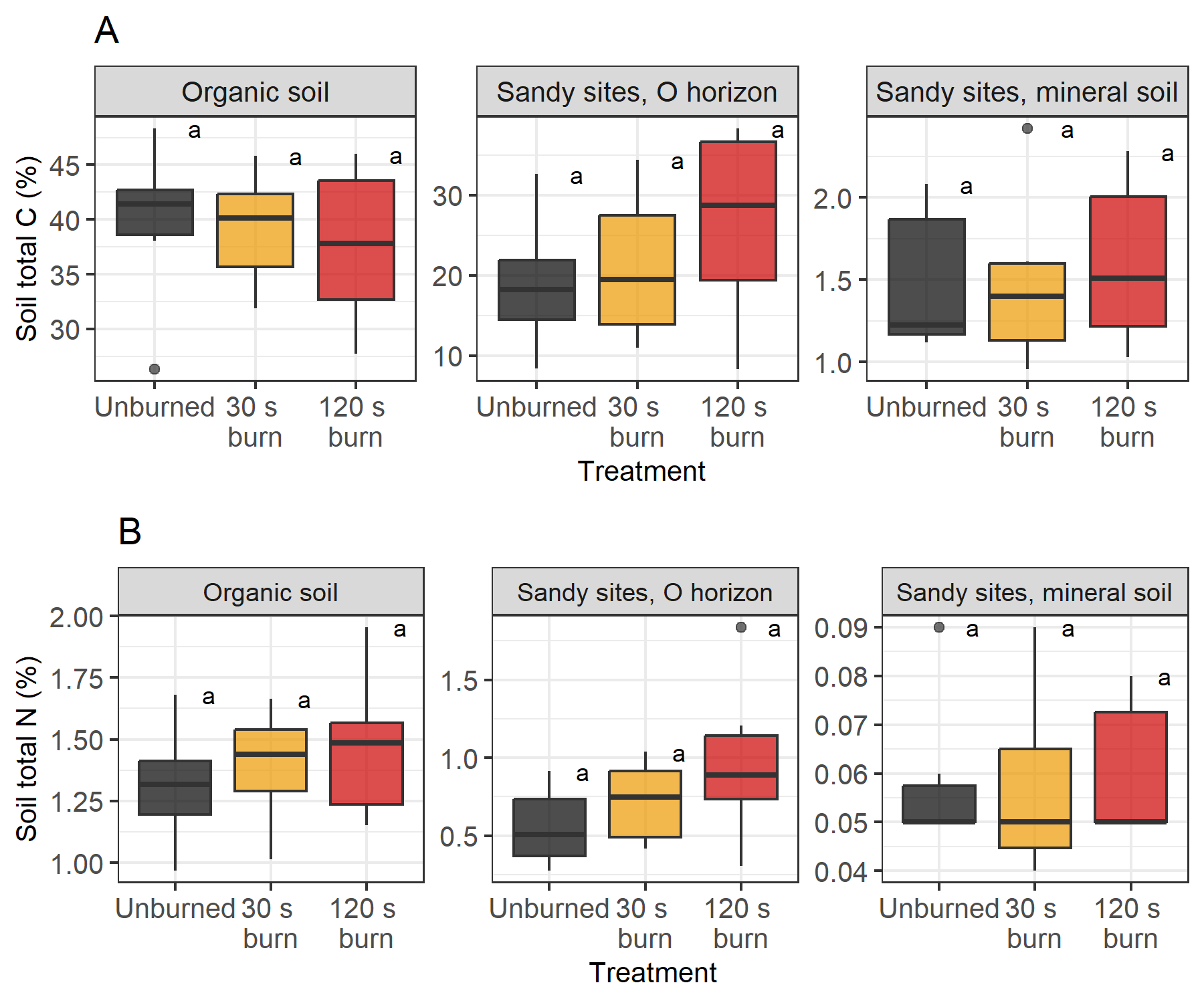

Supplementary Figure 7. Soil (A) C and (B) N concentrations with heat dose treatment in organic soil (left panels), sandy O horizons (middle panels), and sandy mineral soil (right panels). Different letters represent statistically significant differences in treatments based on ANOVA and Tukey’s HSD (p<0.05). The central horizontal line indicates the median, the upper and lower bounds of the box indicate the inter-quartile range (IQR), the upper and lower whiskers reach the largest or smallest values within a maximum of 1.5 * IQR, and data beyond the whiskers are indicated as individual points.

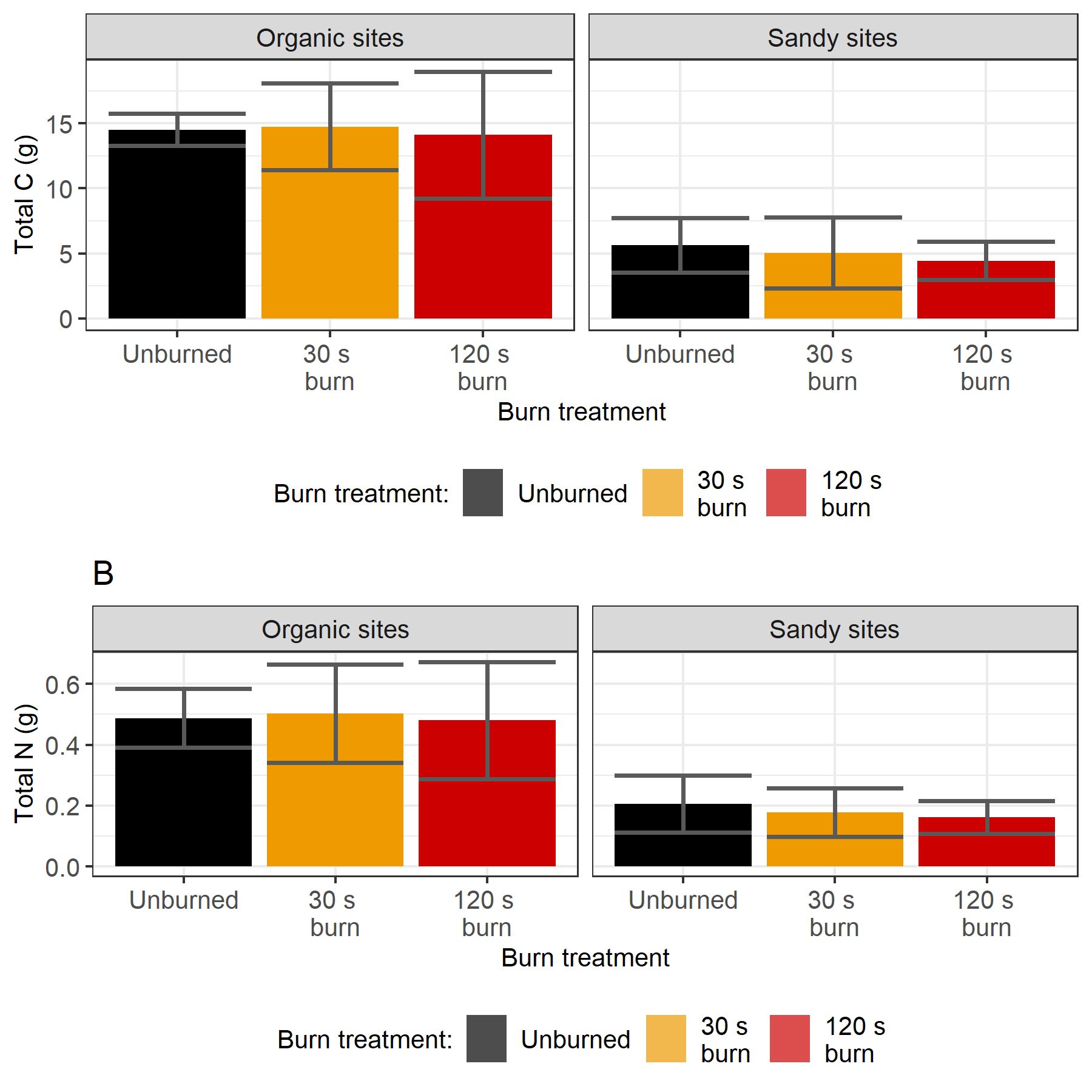

Supplementary Figure 8. Estimates of C mass (top) and N mass (bottom) in organic cores (left panel) and sandy cores (right panel). Error bars represent one standard deviation from the mean.

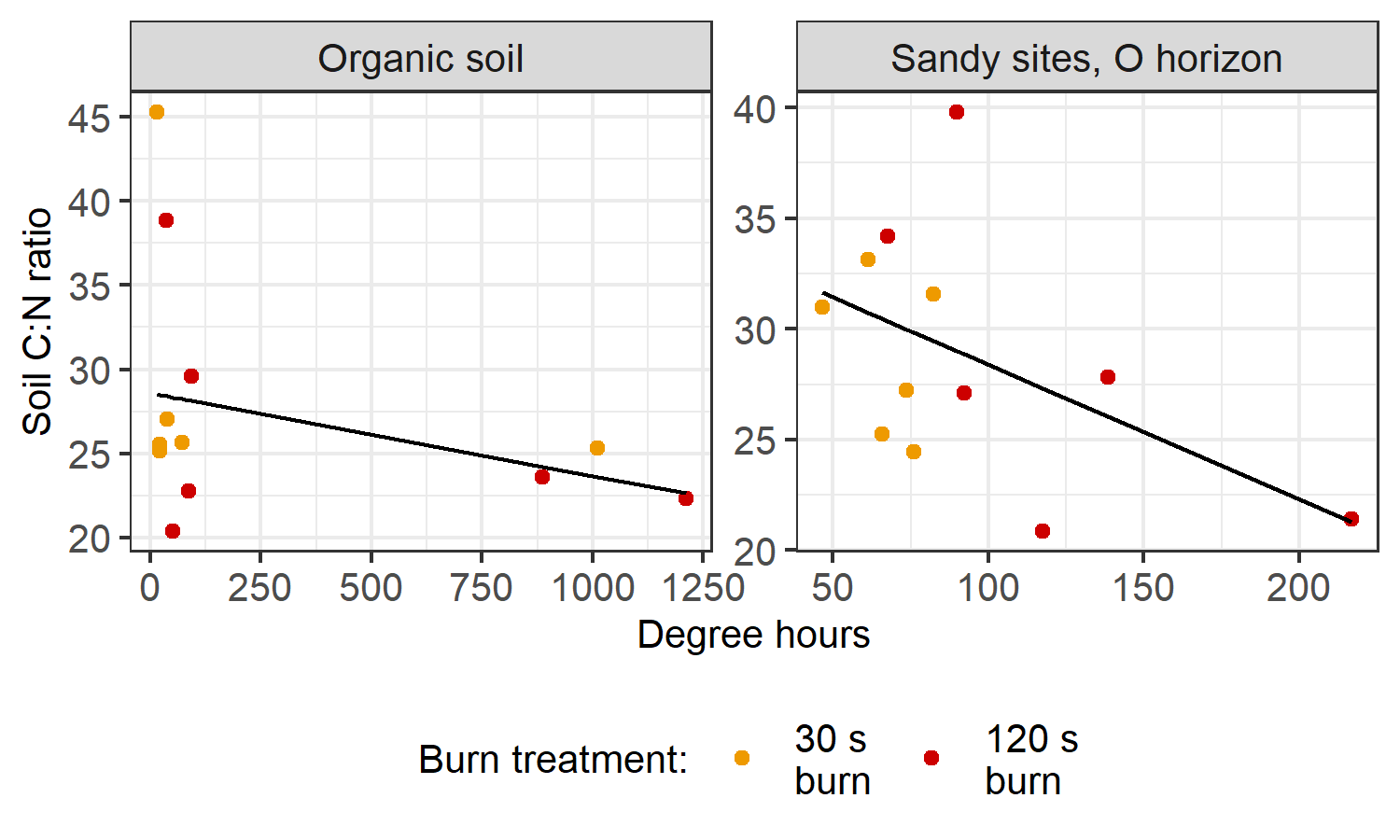

Supplementary Figure 9. Soil C:N ratios two days post-burn vs. degree hours (DH) in organic soils (left panel, y = 28.586 + -0.005x) and O horizons of sandy soils (right panel, y = 34.469 + -0.061x) (n=12 soil horizon samples, p=0.04, $R_{adj.}^{2}$=0.10). DH were calculated based on thermocouple placed at O-mineral interface or at core midpoint. We used a linear model including an interaction term between site type and DH to test for a significant correlation between DH and soil C:N (interation term between site type and DH, p=0.02).

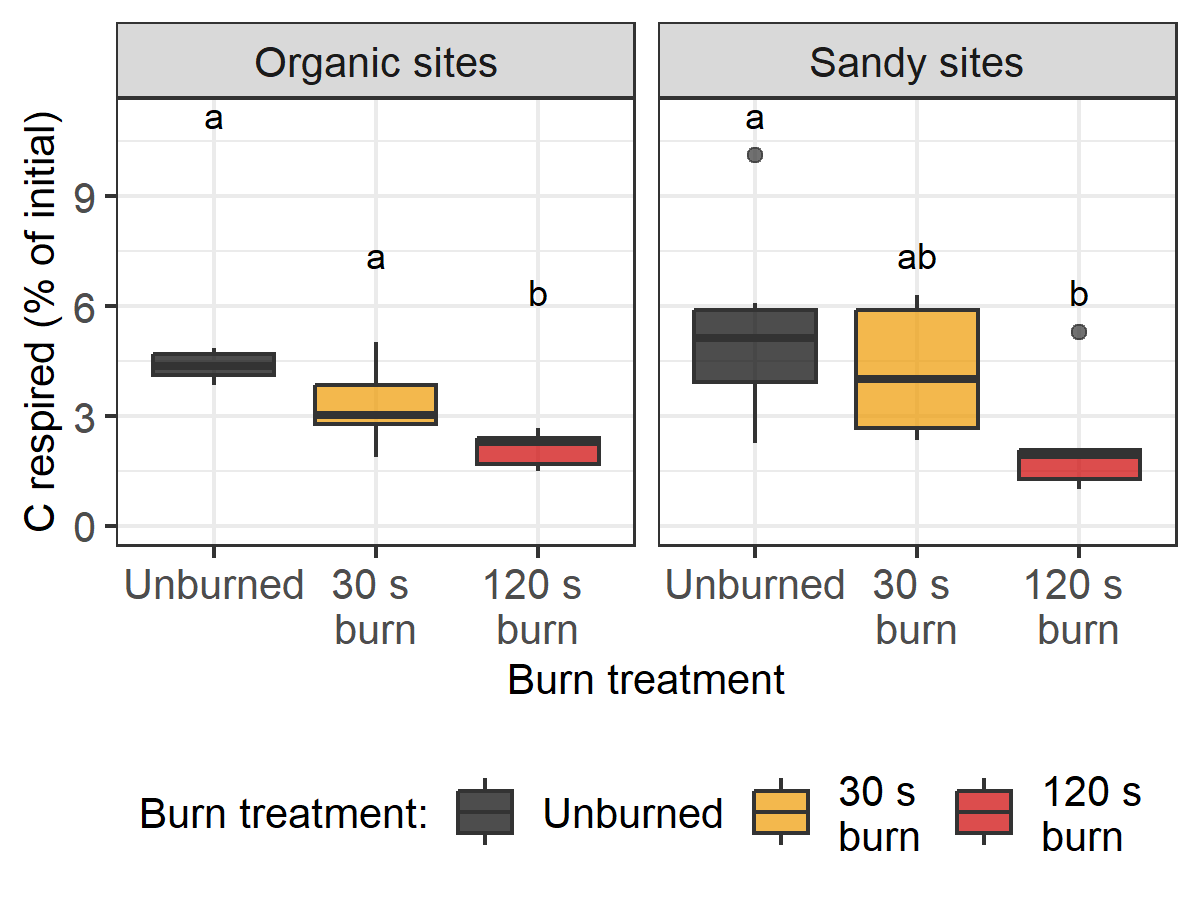

Supplementary Figure 10. C respired (as a percent of post-burn total C) over the entire 70 day incubation of organic cores (left panel) and sandy cores (right panel). Different letters represent statistically significant differences in treatments based on ANOVA and Tukey’s HSD (p<0.05). The central horizontal line indicates the median, the upper and lower bounds of the box indicate the inter-quartile range (IQR), the upper and lower whiskers reach the largest or smallest values within a maximum of 1.5 * IQR, and data beyond the whiskers are indicated as individual points.

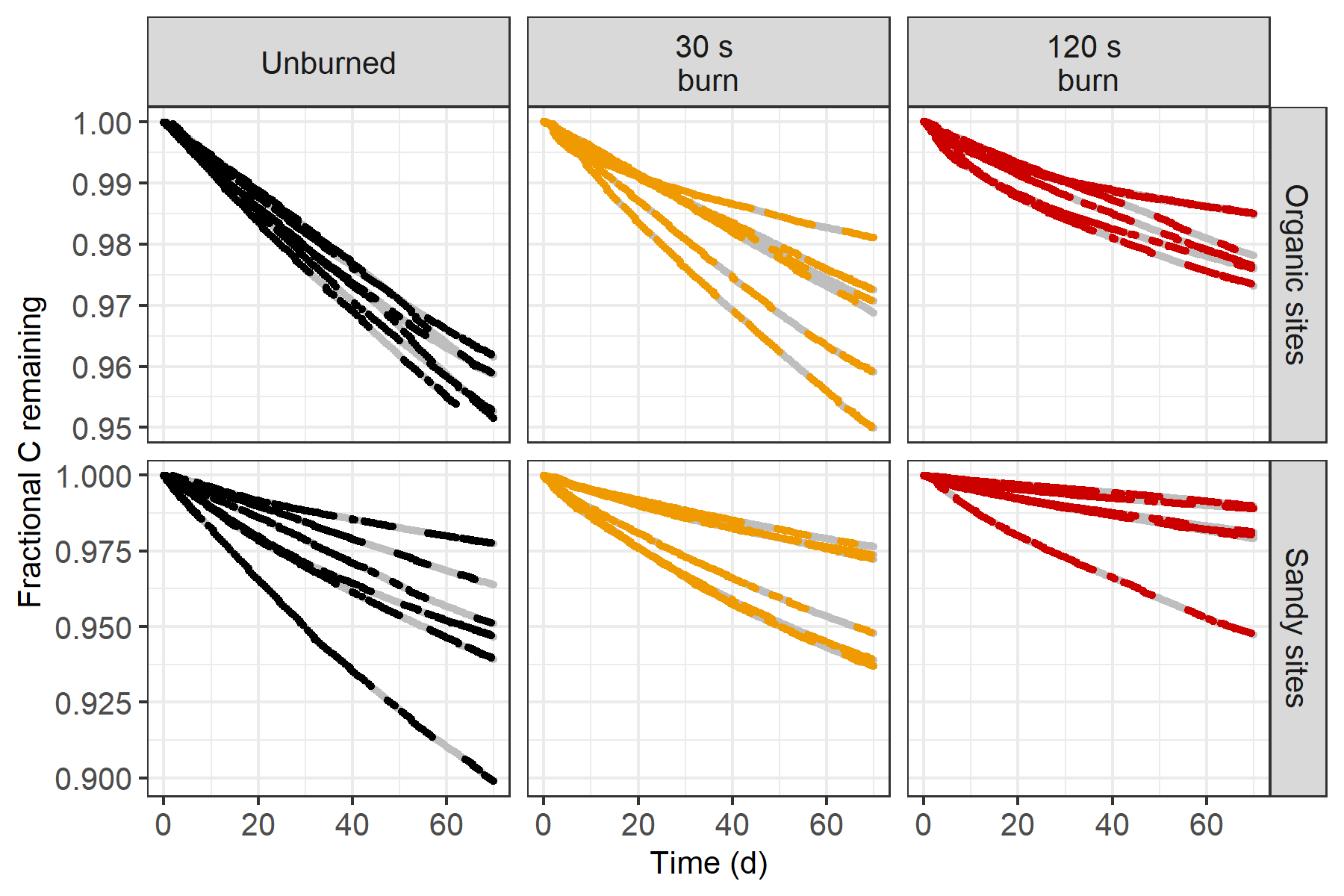

Supplementary Figure 11. Fraction of remaining post-burn total C over the course of a 70-day incubation of organic cores (top panels) and sandy cores (bottom panels). Due to instrument issues, we use average respiration rates to account for periods of missing data (represented here as grey points).

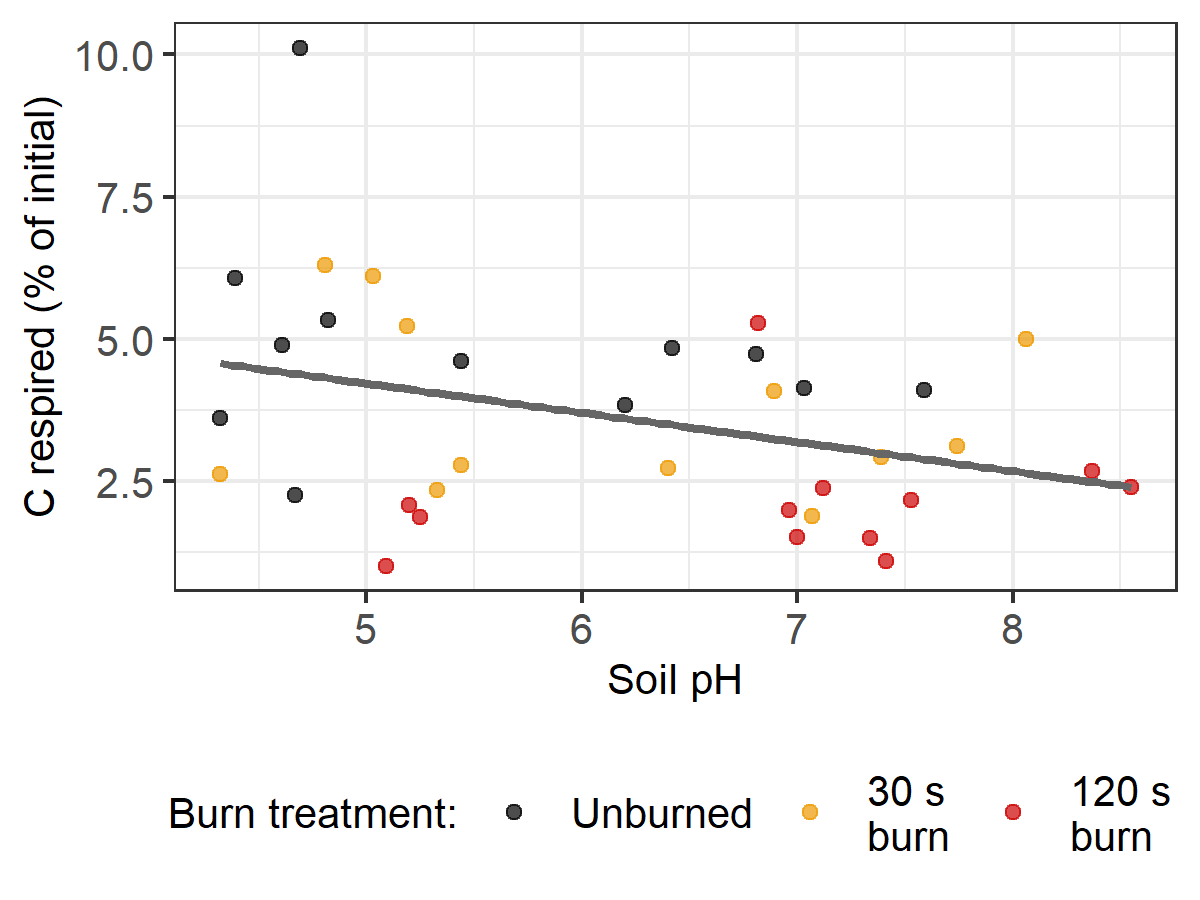

Supplementary Figure 12. C respired (as a percent of post-burn total C) vs. soil pH over the entire 70-day incubation (y = 6.792 - 0.514x; p=0.04, $R_{adj.}^{2}$=0.10).

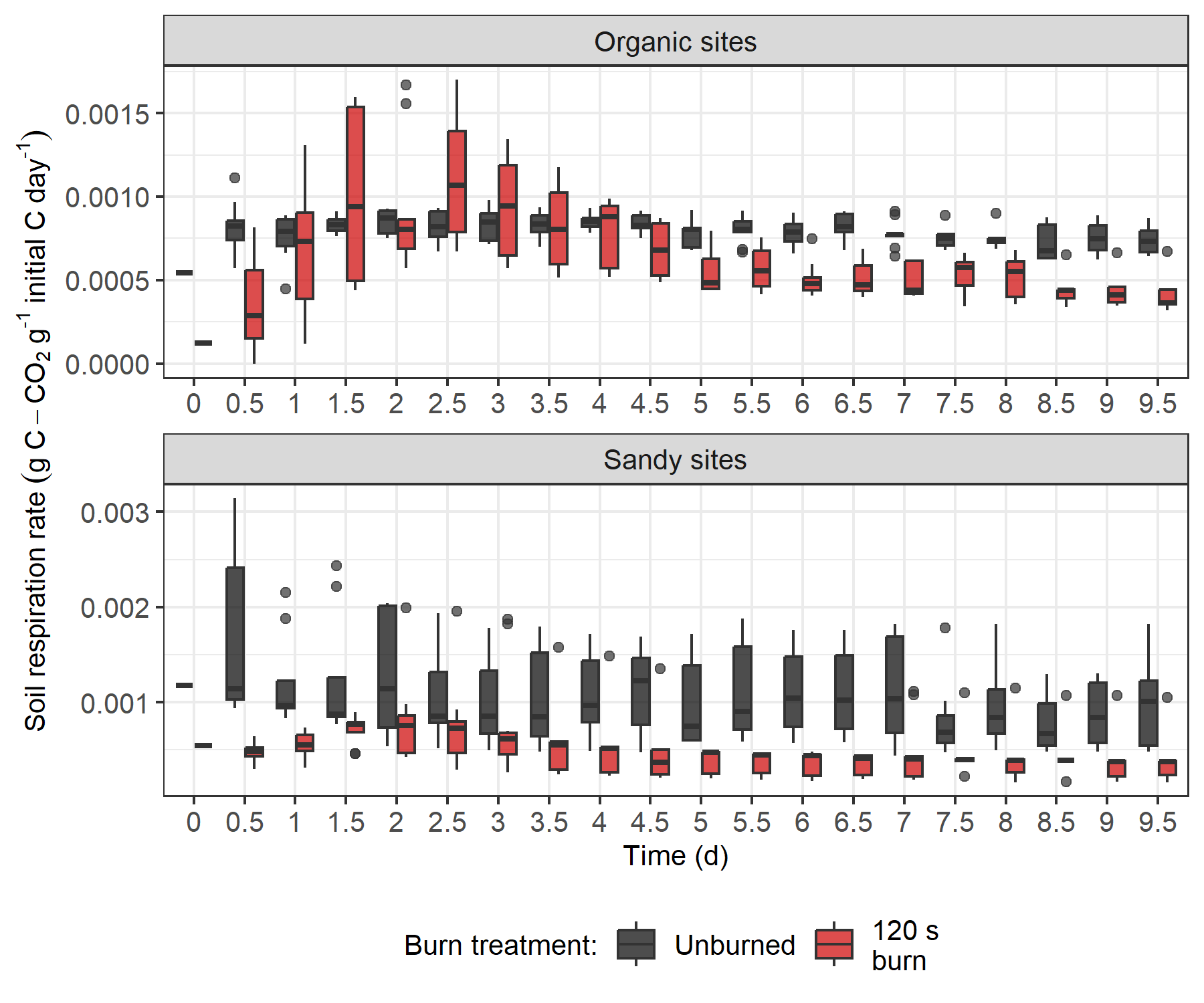

Supplementary Figure 13. Soil respiration rates for the first 10 days of the 70-day incubation of organic soil (top panel) and sandy soil (bottom panel). Initial C refers to the total C at the beginning of the incubation. The central horizontal line indicates the median, the upper and lower bounds of the box indicate the inter-quartile range (IQR), the upper and lower whiskers reach the largest or smallest values within a maximum of 1.5 * IQR, and data beyond the whiskers are indicated as individual points.
